# Sex-Dimorphic Neural Memory Shapes Pancreatic Tissue Resilience

**DOI:** 10.64898/2026.06.15.732370

**Authors:** Rute MM Ferreira, Claudio Ballabio, Ernesto Rodriguez, Adam Karoutas, Probir Chrakavarti, Enrica Martinelli, Marco Stazi, Sara Salgueiro Torres, Victoria Bridgeman, Stefanie Ruhland, Leanne Li, James N. Sleigh, Ilaria Malanchi

## Abstract

Epithelial cells can encode prior damage into lasting epigenetic and functional states, enabling a primed response to future insults. In the pancreas, acute injury induces reversible acinar cell reprogramming toward a progenitor-like identity that persists beyond repair, supporting resilience to recurrent injury but creating a permissive state for malignant transformation. Given the central role of the tissue niche in stem cell regulation, we investigated microenvironmental adaptations that sustain this primed epithelial state. Using genetic mouse models and ex vivo organoid co-cultures, we identify a sex-specific sensory neural memory after pancreatitis that sustains long-term epithelial plasticity through a CGRP-dependent neuron–epithelial axis. We show that sex differences in acute inflammation drive neutrophil-dependent suppression of neural activation in females, decoupling neural memory from epithelial plasticity after repair. In males, neural memory promotes post-injury plasticity, revealing tissue memory as coordinated adaptation between epithelial progenitors and their niche.

## INTRODUCTION

Over the past decade, our understanding of how tissues acquire resilience to repeated insults has evolved substantially, leading to the emergence of the concept of epithelial injury memory as an epigenetically encoded adaptive program. The capacity to “remember” prior insults and mount an accelerated response to subsequent injury, once thought to be exclusive to the adaptive immune system, is now recognised as a characteristic feature of epithelial cells as well. Through this process, epithelial tissues acquire the ability to respond more efficiently and regenerate more rapidly following recurrent injury or inflammation^1^. In the pancreas, activation of epithelial plasticity during pancreatitis, which enables acinar cell dedifferentiation and tissue repair, has been shown to induce an epigenetic state that promotes carcinogenesis^2^. Moreover, the inflammatory response associated with pancreatitis was shown to establish a persistent inflammatory epigenetic program in acinar cells that remains long after tissue repair. While this state enables a faster regenerative response upon secondary injury, it also maintains the tissue in a condition that is more susceptible to oncogenic transformation^3,4^.

While the concept of inflammatory epithelial memory appears to reflect an intrinsic property of epithelial cells, whereby cells maintain a less differentiated state conducive to both tissue repair and carcinogenesis, it remains unclear whether, within the tissue, this phenotype is self-sustained or, like stemness-associated features of tissue progenitors, requires active maintenance by the surrounding tissue microenvironment. If the latter were true, the memory of the injury within the tissue would need to co-exist across the different cellular compartments, the epithelial progenitor compartment, as well as in cellular constituents of the niche.

In this study, we investigated pancreatic tissue memory of injury in mature adult male and female mice to determine whether cellular compartments, beyond the epithelium, display persistent activity following tissue repair. Here, we show that nociceptor neuron activity, initiated during pancreatic injury and active component of the tissue repair response, persists long after tissue regeneration has been completed. Furthermore, we demonstrate that this neural memory of injury is specific to male mice, where it is required to sustain the epithelial memory necessary for tissue resilience upon secondary insult.

## RESULTS

### Sex dimorphism in sustained post-injury pancreatic molecular remodelling

Mature organs maintain high tissue integrity despite emerging age-associated declines in regenerative capacity compared to young organs and represent a stage in which tissue resilience mechanisms may may become increasingly critical to support long-term homeostasis. As epithelial memory maintenance is a key feature impacting both pancreatic resilience to injury and cancer predisposition, we investigated regulators of epithelial memory of injury in mature pancreas of 6 months old mice. We employed a reversible model of pancreatic injury, acute pancreatitis (AP), in adult male and female mice and assessed pancreatic remodelling upon injury induction. Acute pancreatitis was achieved by administration of supraphysiological doses of the cholecystokinin analogue caerulein, over two days, which leads to an overactivation of CCK1 receptors on pancreatic acinar cells and consequent basolareral release of secretory granules into the pancreatic parenchyma. This triggers acute inflammation upon injury and a rapid transcriptional shutdown of acinar cell function, de-differentiation and activation of an acinar to ductal metaplasia (ADM)^5^. ADM characterises the inflammatory phase of the injury, while their re-differentiation into acinar cells allow tissue repair and resolution of inflammation one week post-damage (Figure1A, 1B and S1A).

Mature pancreas of male and female animals were equally responsive to caerulein administration exhibiting comparable release of intracellular pancreatic amylase 24 hours post-AP (Figure 1C and S1B). There was also no differences in upregulation of the injury and progenitor marker STMN1^6,7^ (Figure 1D).

**Figure 1.**
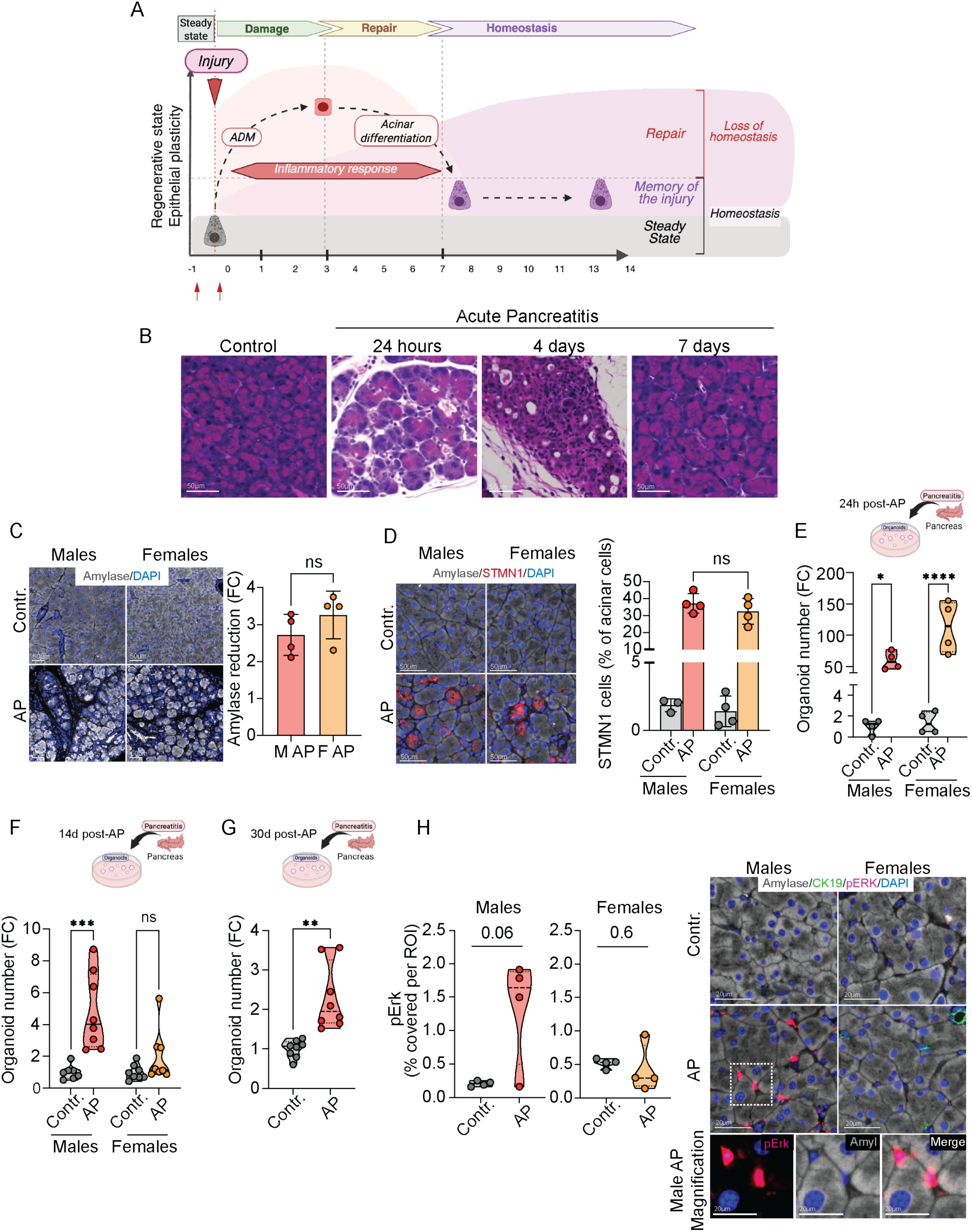
Sex-specific retention of AP-induced pancreatic epithelial plasticity. **(A)** Schematic of epithelial plasticity kinetics during acute pancreatitis (AP) and repair, illustrating low baseline plasticity at steady state, peak induction during acinar-to-ductal metaplasia (ADM), partial resolution upon acinar re-differentiation, and persistently elevated plasticity at post-repair heterostasis (injury memory). **(B)** Representative H&E images of male pancreas from control and AP at 24 hours, 4 days, and 7 days after the final caerulein injection, illustrating histological progression from injury to resolution. Scale bars are indicted in the figures. **(C)** Representative amylase/DAPI immunofluorescence images (left) and quantification (right) of pancreatic parenchyma area with amylase loss in male and female AP mice at 24 hours post-injury (Male AP n=4, Female AP n=4). Data are represented as mean ± SD (fold change relative to sex-matched control). Scale bars are indicted in the figures. **(D)** Representative amylase/Stathmin1 (STMN1)/DAPI immunofluorescence images (left) and quantification (right) of STMN1+ acinar cells in control and AP male and female pancreas at 24 hours after final caerulein injection (Male control n=3, Male AP n=4, Female control n=4, Female AP n=4). Data are represented as mean ± SD (fold change relative to sex-matched control). Scale bars are indicted in the figures. **(E–G)** Schematics (top) and organoid formation efficiency (bottom; number of organoids formed per well, normalised to sex-matched control mean) from male and female mice at 24 hours (E; Male control n=4, Male AP n=4, Female control n=4, Female AP n=4), 14 days (F; Male control n=7, Male AP n=8, Female control n=8, Female AP n=8), and 30 days post AP (G; males only; Male control n=8, Male AP n=8). Data are represented as median (violin plots). t-test. **(H)** Quantification (left) of pancreatic parenchyma area covered by pErk signal within ROIs and representative amylase/CK19/pErk/DAPI immunofluorescence images (right) from male and female control and AP mice 14 days post-AP (Male control n=4, Male AP n=4, Female control n=4, Female AP n=4). Data are represented as median (violin plots). Scale bars are indicted in the figures. *p < 0.05, **p < 0.01, *** p < 0.001 by Unpaired Welch’s t-test See also Figure S1.

Epithelial plasticity enables cells to dynamically adjust their identity, behaviour, and regenerative potential in response to injury, therefore increased epithelial plasticity serves as a key mechanism through which epithelial progenitors acquire an activated regenerative state. Therefore, we refer to the level of epithelial plasticity as a measure of the injury-induced regenerative activation of epithelial progenitors. As epithelial plasticity is also needed to initiate and sustain self-renewing organoid structures ex vivo, the level of pancreatic regenerative potential associated with ADM can be evaluated by organoid formation assays^8^. Here, the number of organoids generated from freshly harvested pancreatic epithelial cells reflects the proportion of cells with the tissue with the level of epithelial plasticity required for organoid initiation at the time of collection. At 24 hours post-AP, the time of acinar dedifferentiation, both male and female pancreatic epithelial cells exhibit extremely high plasticity, forming significantly more organoids, as compared to matched controls (Figure 1E and S1C). As the epigenetic state of epithelial cells was reported to remain with a higher level of regenerative plasticity after the injury^3^, we tested whether this epithelial memory could also be detected using organoid formation assays after tissue repair. Interestingly, while pancreatic epithelial cells from male animals retained increased organoid formation capacity at 7, 14 and 30 days after AP, pancreatic epithelial cells from female animals did not display this feature post-AP (Figure 1E-G and S1D), suggesting that pancreas from mature male mice display an enhanced long-term increased in epithelial plasticity compared to female mice that can be efficiently detected in organoid assays.

We investigated further the differences in molecular adaptations retained upon tissue repair in the post injury state of male and female pancreatic epithelia. Previous studies have shown that Erk phosphorylation (pErk) remains elevated in pancreatic lysates up to 12 weeks post-injury^4^. At 14 days post injury, immunofluorescence analysis of pancreas of female and male animals demonstrated reinforced the sex dimorphism, with post-injury male, but not female pancreas containing more pErk-positive cells compared to controls (Figure 1H).

These findings uncover a sex-dimorphic heterostasis. While both sexes respond similarly in the early stages of injury, males retain higher levels of plasticity after repair compared to females.

### Sex dimorphism in neuronal plasticity in response to AP

While pErk signalling has been implicated in the maintenance of the tissue memory of pancreatic injury^4^, the cells in which this signalling is present has not been addressed. Surprisingly, we observed that pErk cells were commonly found in the periacinar interstitium (Figure 1H, higher magnification). These interstitial spaces between acini, within lobules, contain many stromal cells and nerve axons^9^. Co-immunofluorescence for pErk showed close proximity with nerve axons (βIIITubulin) compared to fibroblasts (aSMA), another abundant component of this periacinal space (Figure 2A).

**Figure 2.**
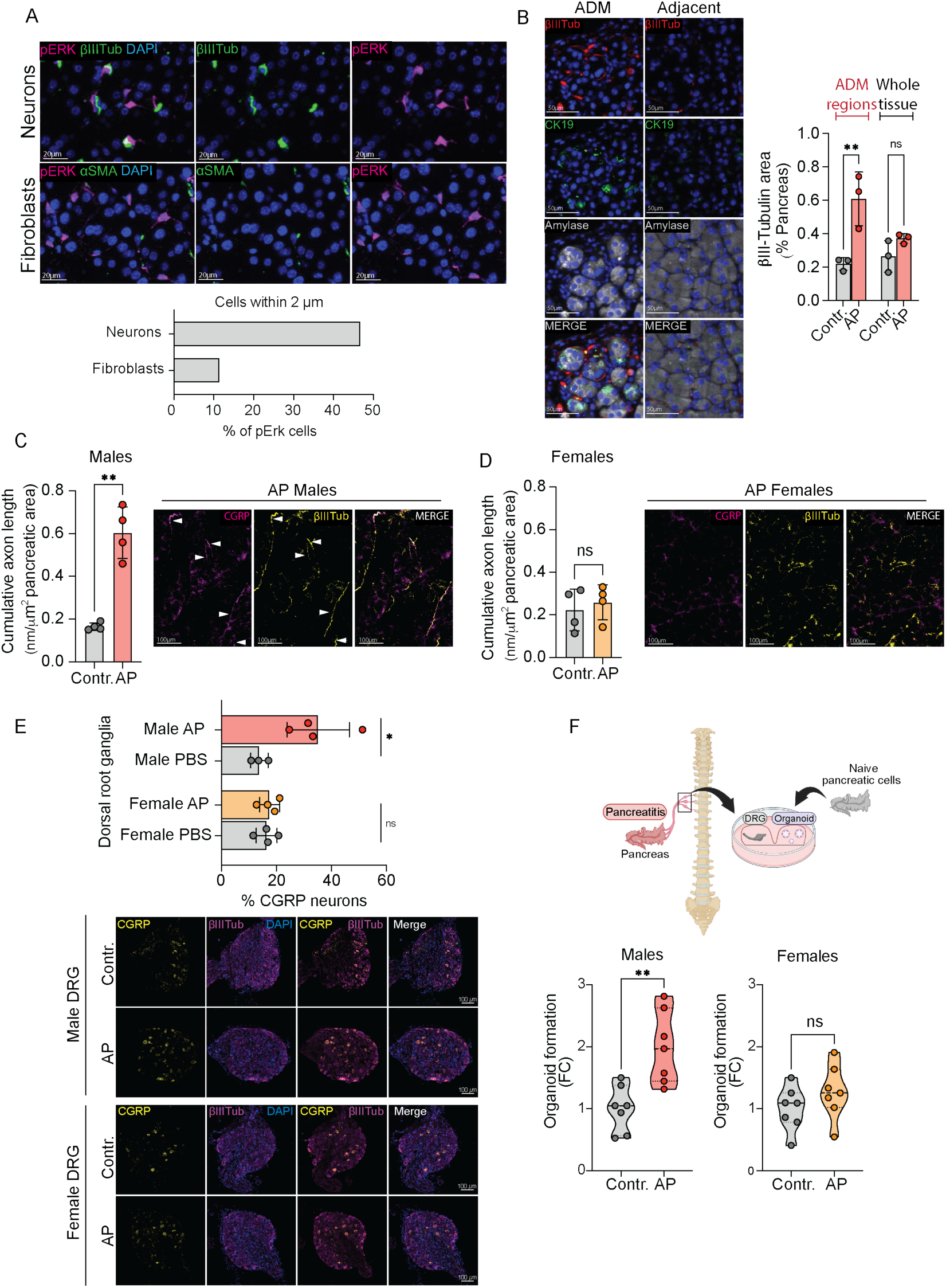
Males and females share structural remodelling but diverge in functional injury-induced peripheral neural plasticity. **(A)** Representative dual immunofluorescence images for pErk/âIII-Tubulin (âIII-Tub; pan-neuronal) and pErk/áSMA (fibroblasts) in male pancreas 14 days post-AP (top), and quantification (bottom) of pErk+ cells within a 2 μm radius of neurons (βIII-Tub) or fibroblasts (αSMA), expressed as percentage of total pErk+ cells (n=230 for βIII-Tub; n=350 for αSMA; n= total pErk+ cells). Scale bars are indicted in the figures. **(B)** Representative amylase/CK19/âIII-Tub/DAPI immunofluorescence images in male AP pancreas 24 hours post-AP showing innervation within ADM and adjacent areas (left), and quantification (right) of βIII-Tub+ area in ADM versus whole pancreas relative to matched control regions (Male control n=3, Male AP n=3). Data are represented as mean ± SD. Scale bars are indicted in the figures. **(C, D)** Cumulative CGRP+ axon length (nm per μm² parenchyma) and representative CGRP/βIII-Tub immunofluorescence images of 100 μm pancreatic sections 24 hours post-AP in males (C; Male control n=4, Male AP n=4) and females (D; Female control n=4, Female AP n=4). Data are represented as mean ± SD. Scale bars are indicted in the figures. **(E)** Quantification (top) of CGRP+ neurons as a percentage of total neurons in pancreas-innervating dorsal root ganglia (DRGs) from male and female mice 24 hours post-AP (Male control n=3, Male AP n=4, Female control n=4, Female AP n=4), and representative CGRP/βIII-Tub/DAPI immunofluorescence images (bottom). Data are represented as mean ± SD. Scale bars are indicted in the figures. **(F)** Schematic (top) of the pancreas-innervating DRGs (pDRGs) and naïve epithelial cell co-culture assay to assess neuron activation and support of epithelial plasticity. pDRGs isolated 24 hours post-AP were co-cultured with naïve pancreatic epithelial cells in organoid formation condition. Organoid formation efficiency (bottom) by sex (Male control n=7, Male AP n=7, Female control n=7, Female AP n=7). Data normalised to sex-matched control mean per experiment. Data are represented as median (violin plots). *p < 0.05, **p < 0.01 by Unpaired Welch’s t-test See also Figure S2.

The close spatial association led us to investigate the structure and function of the intrapancreatic neural network in the context of AP. We performed whole organ 3D imaging for the pan-neuronal marker βIIITubulin on pancreas from control and AP mice 24 hours post-AP onset. As previously described^10^, control pancreas exhibited an organised and hierarchical network of βIIITubulin nerve fibres running along the ductal tree (Figure S2A). Pancreas isolated 24 hours post-AP displayed significant structural changes exhibiting a disorganised pattern, with nerve fibres concentrated in discrete foci throughout the parenchyma (Figure S2A). We performed 2D immunofluorescence to quantify this neural remodelling in control and AP pancreas and observed that, as soon as 24 hours post-AP onset, there was a significant increase in nerve fibres density specifically in areas of damage, ADM regions, identified by co-localisation of CK19 and amylase on acinar cells (Figure 2B). This neural structural remodelling induced by AP did not differ between males and females (Figure S2B).

As neural changes in pancreatic disease extend beyond alterations in fibre density^11^, we next sought to define the compositional remodelling of intrapancreatic innervation in response to AP. Of the three branches of the PNS innervating the pancreas, sensory afferents are the predominant neural drivers of pancreatitis onset and progression^12^. Amongst the neuropeptides released by sensory afferents, CGRP represents one of the principal mediators of their effects on pancreatic tissue^11^. Immunohistochemistry analysis of CGRP fibres in pancreatic tissues 24 hours post-AP onset revealed that AP induces a significant increase in cumulative CGRP^+^ fibre length exclusively in males (Figure 2C and 2D). Similarly, pancreas-innervating dorsal root ganglia (pDRG), where sensory soma reside, of male but not female animals exhibited a significant increase in the proportion of CGRP^+^ neurons 24 hour post-AP induction (Figure 2E).

Given the sex-dimorphic neurochemical remodelling, we hypothesised that AP may drive sex-dependent changes in the functional engagement of pDRG neurons. In AP, sensory neurons are known to regulate the extent of pancreatitis by modulating the immune and endothelial pancreatic compartment^12^. Since there were no differences in AP onset between male and female animals, we reasoned that such changes could be reflected in DRG regulation of the epithelia after the injury. Little is known of the function of sensory neuron activation in epithelial biology, however, studies in pancreatic cancer have shown that cancer cells co-cultured with DRGs have enhanced proliferative capacity and upregulation of pathways associated with plasticity^13^. Hence, we tested whether pDRG explants enhance epithelial plasticity to form organoids of naïve pancreatic epithelial cells. To this end, we performed organoid co-cultures of pDRGs isolated from control and AP pancreas with naïve pancreatic primary epithelial cells (Figure 2F, top panel). Aligned with a higher sensory engagement, male pDRG robustly enhanced naïve pancreatic epithelial plasticity, while female pDRGs did not show this activity (Figure 2F).

Collectively, these findings indicate that AP induces rapid structural remodelling of the intrapancreatic neural network in both sexes. However, it drives a sex-dimorphic functional adaptation whereby males exhibit a heightened sensory activity, which is defined as the ability to boost epithelial plasticity *ex vivo* and is reflected in an increased CGRP-positive innervation in pancreatic tissue.

### Male-specific nociceptor dependency of persistent epithelial plasticity following pancreatitis

A body of literature have shown the engagement of sensory neuron at the time of injury and their key role in regulating inflammation during the phase of pancreatitis^12^. Activation of pancreatic sensory neurons can promote neurogenic inflammation through the release of neurotransmitter (substance P and CGRP), leading to vasodilation, plasma extravasation, and pain, and may thereby amplify local inflammation during acute pancreatitis^14^. To confirm that the described neural activity to support epithelial plasticity upon AP is gained by DRG neurons stimulated directly by local injury/inflammatory signals within the tissue, we compared the ability of male thoracic level 10 (T10) pDRGs, which innervate the injured pancreas, and lumbar level 1 (L1) DRGs, which innervate the colon lying outside the primary splanchnic innervation territory of the pancreas^15^. L1 DRGs isolated 24 hours from the same mice post-AP failed to support pancreatic epithelial plasticity *ex vivo* following AP, pointing at pancreas-specific neural response (Figure 3A).

**Figure 3.**
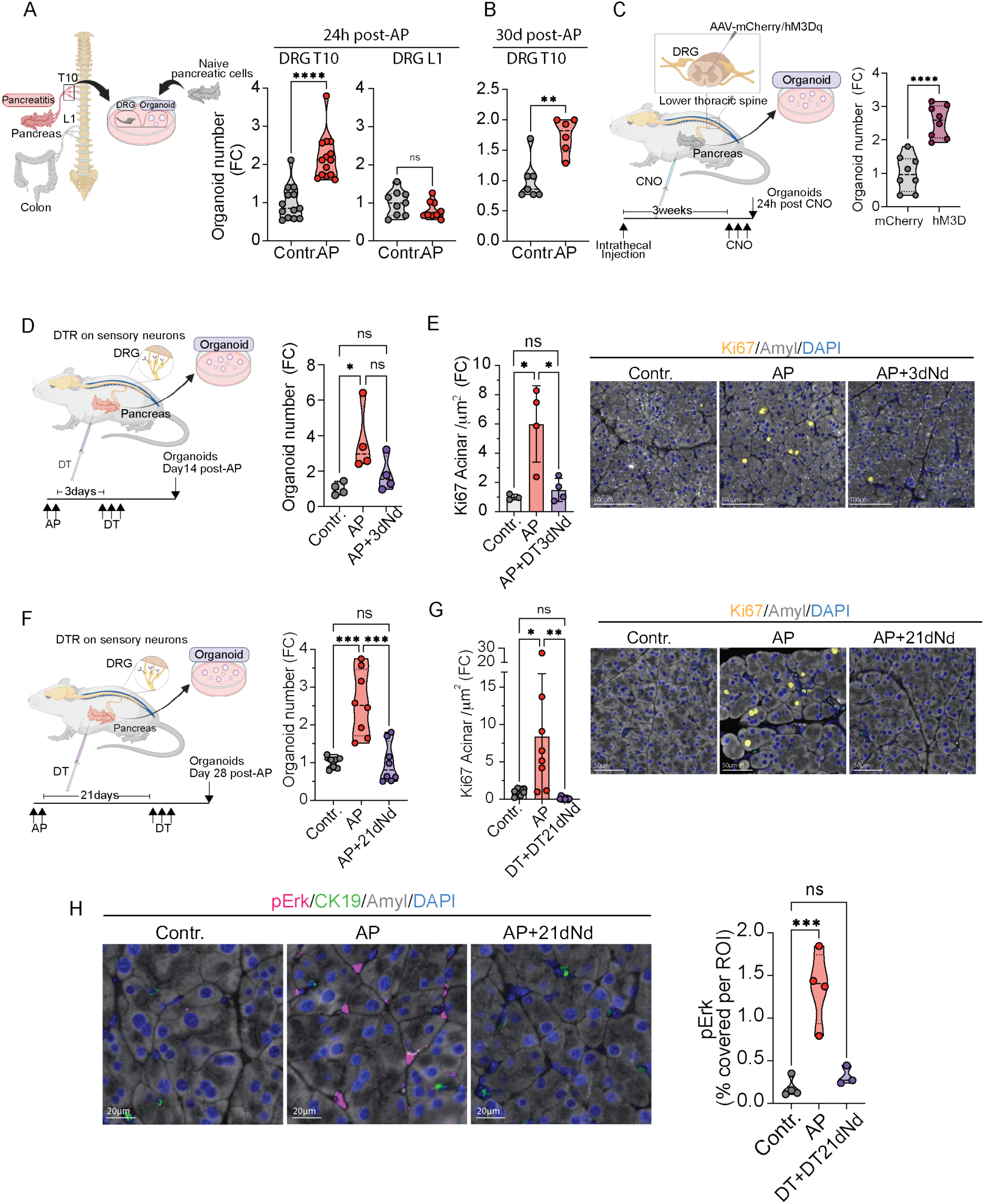
Acute pancreatitis activates nociceptor neurons in males to support post-injury retention of epithelial memory. **(A)** Schematic (left) of pDRG co-culture assay for panels A and B. Pancreatic (T10)-and colon (L1)-innervating DRGs from male mice 24 hours post-AP were co-cultured with naïve pancreatic epithelial cells to assess plasticity support by organoid formation efficiency. Organoid formation efficiency (right) of naïve epithelial cells in the presence of DRGs within innervation territory T10 and L1 (T10: Male control n=13, AP n=14; L1: Male control n=9, AP n=9). Data normalised to control mean per experiment. Data are represented as median (violin plots). **(B)** Organoid formation efficiency from male pDRG and naïve epithelial cell co-cultures, as in A, 30 days post-AP (Male control n=7, AP n=6). Data normalised to control mean per experiment. Data are represented as median (violin plots). **(C)** Schematic (left) of DREADD-mediated sensory neuron activation. hM3Dq or mCherry (control) was expressed in pancreas-territory sensory neurons by intrathecal viral injection; pancreas was collected 24 hours after CNO administration. Organoid formation efficiency (right; 4 technical replicates per mouse, n=2 mice per group). Data normalised to mCherry control. Data are represented as median (violin plots). **(D, E)** Early nociceptor depletion experiment. Schematic (D, left): AP was induced in Scn10a-Cre;R26-DTR males; nociceptor neurons were depleted by diphtheria toxin (DT) 3 days post-AP, with analysis at day 14. Organoid formation efficiency (D, right; Contr. n=4, AP n=4, AP+3dNd n=4). Data normalised to control mean. Data are represented as median (violin plots); one-way ANOVA. Representative amylase/Ki67/DAPI immunofluorescence images (E, left) and quantification of Ki67+ acinar cell density per μm² pancreatic parenchyma (E, right) from the same animals. Data are represented as mean ± SD. Scale bars are indicted in the figures. **(F–H)** Late nociceptor depletion experiment. Schematic (F, left): nociceptor neurons were depleted by DT 21 days post-AP in Scn10a-Cre;R26-DTR males, with analysis at day 28. Organoid formation efficiency (F, right; Contr. n=7, AP n=8, AP+21dNd n=8). Data normalised to control mean per experiment. Data are represented as median (violin plots); one-way ANOVA. Ki67+ acinar cell density per μm² parenchyma (G, left) and representative amylase/Ki67/DAPI immunofluorescence images (G, right; Contr. n=8, AP n=8, AP+21dNd n=8). Data are represented as mean ± SD; one-way ANOVA. Representative amylase/CK19/pErk/DAPI immunofluorescence images (H, left) and pErk+ parenchyma area within ROIs (H, right; Contr. n=4, AP n=4, AP+21dNd n=3). Data are represented as median (violin plots); Scale bars are indicted in the figures. *p < 0.05, **p < 0.01, *** p < 0.001, **** p < 0.0001 by Unpaired Welch’s t-test (A-C) and one-way ANOVA (D-H) See also Figure S3.

We next assess, whether this activation persists upon resolution of inflammation, male pDRGs which would have gain this activity during tissue injury, were isolated at defined timepoints following resolution of inflammation and co-cultured with naïve pancreatic epithelial cells. Strikingly, pDRGs retained the capacity to enhance epithelial plasticity for at least one month after AP resolution (Figure 3B and S3A).

These results suggest that the AP-induced pancreatic environment in male mice drives persistent sensory neuron activity to enhance epithelial plasticity in *ex vivo* co-cultures. To test if this ability to enhance epithelial plasticity upon neural activation could be detected within the tissue, we assessed whether directly nociceptor activation outside of the inflammation induced by tissue injury was sufficient to enhanced epithelial plasticity. To this, we activated pancreas-innervating nociceptors by delivering a Cre-dependent AAV encoding the excitatory Designer Receptor Exclusively Activated by Designer Drugs (DREADD) receptor hM3Dq intrathecally into Scn10a-Cre mice. Binding of a synthetic ligand, such as Clozapine N-oxide (CNO), leads to selective activation of DREADD-expressing neurons^16^. Virus delivery to lower spinal levels of Scn10a-Cre mice enabled the conditional and selective activation of nociceptor DRG within the primary innervation territory of the pancreas. Pancreatic epithelial cells from sensory activated mice exhibited significantly higher organoid-forming efficiency compared cells from control animals (Figure 3C). This evidence demonstrate that nociceptor activation alone is sufficient to enhance pancreatic epithelial plasticity in the tissue and establishes a direct causal link between sensory neuron activity and epithelial regenerative capacity.

Pancreatic regeneration following AP proceeds through three distinct phases: the acquisition of a progenitor-like state through ADM, the subsequent redifferentiation to acinar identity underscoring repair and inflammation resolution, and finally the retention of a tissue regenerative memory characterised by enhanced epithelial plasticity persisting beyond injury resolution^3,4^. While these three phases are sequentially coupled under normal regenerative conditions, they are governed by distinct molecular programs (Figure 1A). To interrogate the contribution of sensory innervation during specific phases of the pancreatic regenerative response, we conditionally ablated nociceptor neurons via diphtheria toxin (DT) administration in Scn10a-DTR mice at specific time points following AP. Sensory denervation 3 days after the last caerulein injection, a timepoint of established acinar dedifferentiation and high inflammation, led to reduced epithelial plasticity measured at day 14 (Figure 3D) No effect of DT treatment alone on epithelial plasticity was detected (Figure S3B). Nociceptor depletion also reversed the previously described^3^ sustained proliferative signature (Figure 3E) and progenitor marker expression (Figure S3C) associated with the post-injury metastable state. Importantly, sensory denervation did not affect the resolution of ADM or the regenerative replacement of acinar cells, and the regain in pancreatic integrity was comparable in control and mice where sensory fibre were depleted during the inflammatory phase (Figure S3D). This evidence suggests that neural engagement after the injury is maintained as neural-memory and supports the capacity of epithelial cells to sustain progenitor programs after repair but is decoupled from inflammatory modulation once inflammation is established and repair programs are engaged.

Together with the prior data showing that neural support of enhanced epithelial plasticity is maintained long-term after injury resolution, these findings led us to directly assess the contribution of the neural activity on the long-term epithelial memory of injury. We depleted sensory neurons one and three weeks after injury and assessed the pancreatic epithelial plasticity 7 days after depletion. At both timepoints, sensory nerve depletion abrogated the persistent higher state of epithelial plasticity, as evidenced by impaired organoid-forming capacity, reduced Ki67-positive and mitotic pH3-positive acinar cells, and loss of STMN1-positive acinar cells (Figure 3F, 3G, S3E-G). Importantly, this was accompanied by a loss of pErk-positive cells within the pancreatic parenchyma (Figure 3H).

Taken together, these results support the conclusion that male sensory neuron activation upon AP triggers a neural-memory state able to directly support epithelial plasticity that is maintained following injury resolution and required to sustain epithelial memory after injury.

### CGRP stimulation promotes epithelial plasticity by limiting acinar differentiation commitment

We next aimed to dissect the mechanism of neuro-epithelial communication. To test whether direct physical interaction was required for sensory neurons to enhance epithelial plasticity, we generated conditioned media (CM) from pDRGs isolated 24 hours post-AP. This was sufficient to enhance organoid formation efficiency from naïve pancreatic epithelial cells as compared to CM from pDRG control animals, suggesting that plasticity is regulated by a paracrine neuronally-derived soluble factors (Figure 4A).

**Figure 4.**
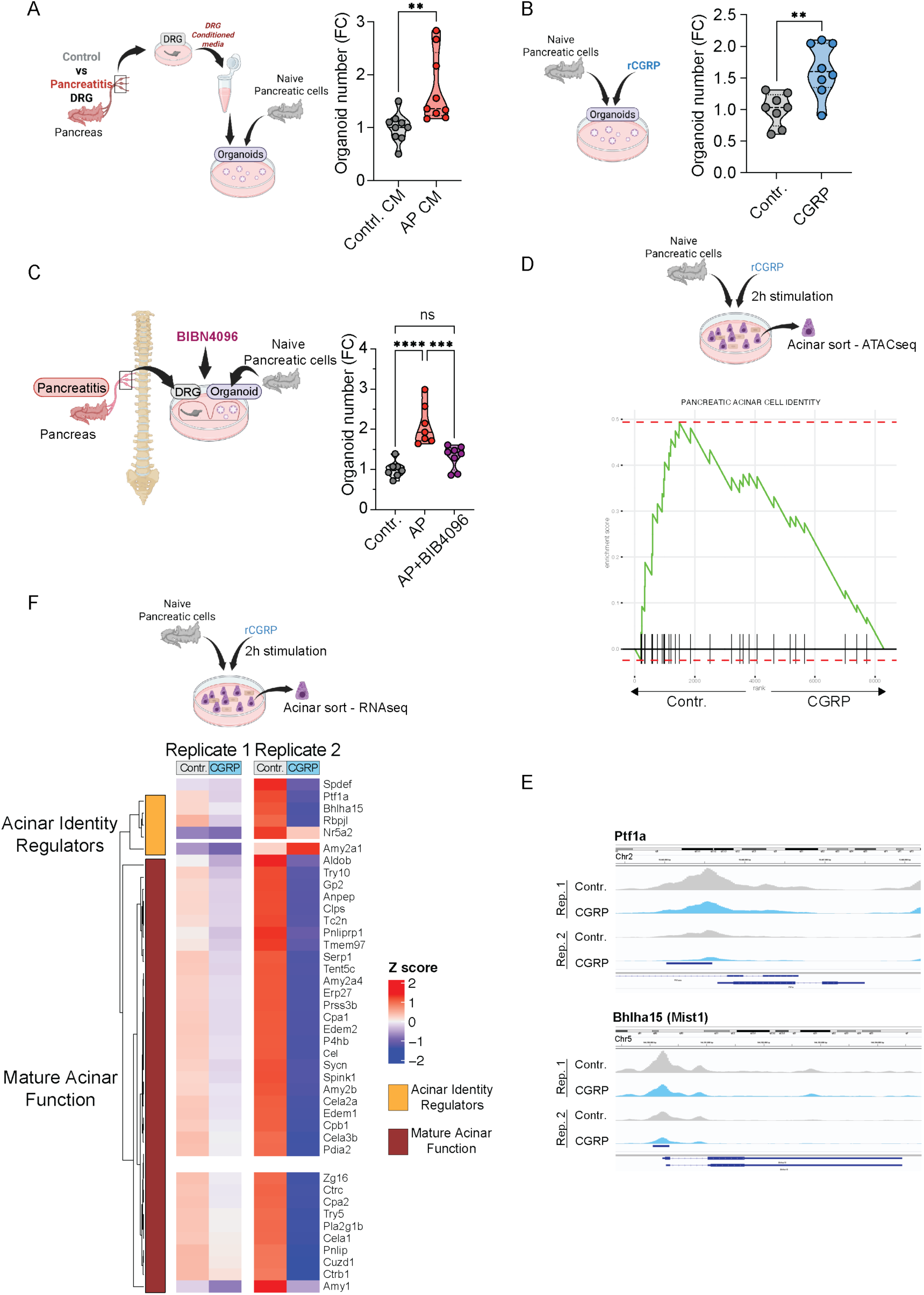
CGRP constrains acinar lineage commitment to preserve epithelial plasticity. **(A)** Schematic (left) of pDRG conditioned media (CM) generation from males 24 hours post-AP applied to naïve epithelial cells in organoid formation assays. Organoid formation efficiency (right; control CM n=9, AP CM n=9). Data normalised to control mean per experiment. Data are represented as median (violin plots). Unpaired Welch’s t-test. **(B)** Schematic (left) of 10 nM CGRP treatment of naïve epithelial cells in organoid formation assays. Organoid formation efficiency (right; control n=8, CGRP n=8). Data normalised to control mean per experiment. Data are represented as median (violin plots). Unpaired Welch’s t-test. **(C)** Schematic (left) of pDRG co-culture assay with CGRP receptor blockade. pDRGs from males 24 hours post-AP were co-cultured with naïve epithelial cells ± 10 μM BIBN-4096 in organoid formation assays. Organoid formation efficiency (right; control n=7, AP n=7, AP+BIBN-4096 n=7). Data normalised to control mean per experiment. Data are represented as median (violin plots). One-way ANOVA. **(D, E)** Schematic (top) of 10 nM CGRP 2 hour stimulation of naïve epithelial cells followed by ATAC-seq of sorted acinar cells. Gene set enrichment plot (D, bottom) showing enrichment of an acinar cell identity signature in control samples. ATAC-seq peaks (E, bottom) at Ptf1a and Bhlha15 loci. **(F)** Schematic (top) of 10 nM CGRP 2 hour stimulation of naïve epithelial cells followed by RNA-seq of sorted acinar cells. Transcriptional profile (bottom) of acinar function and maturation genes in control and CGRP-stimulated cells in paired replicate samples. *p < 0.05, **p < 0.01, *** p < 0.001, **** p < 0.0001 by Unpaired Welch’s t-test (A-B) and one-way ANOVA (C) See also Figure S4.

Given that CGRP-positive fibre density was increased in the male pancreas following AP, we next asked whether CGRP mediates the observed neuro-epithelial support of pancreatic plasticity. Treating naïve pancreatic epithelial cells with synthetic CGRP peptides significantly increased their organoid-forming efficiency (Figure 4B). Conversely, co-culture of naïve pancreatic epithelial cells with post-AP pDRGs in the presence of BIBN-4096, a selective small molecule antagonist of the CLR/RAMP1 CGRP receptor complex^17^, abrogated pDRG-mediated enhancement of organoid-forming capacity (Figure 4C). Importantly, BIBN-4096 did not directly impact epithelial cell plasticity (Figure S4A). These data suggest CGRP stimulation of epithelial cells is important for pDRG-dependent support of epithelial plasticity.

The enhanced epithelial plasticity associated with post-injury tissue memory has been attributed to persistent epigenetic remodelling induced at the time of injury^2–4^. To determine the potential chromatin remodelling caused by direct CGRP stimulation, pancreatic epithelial cells isolated from naïve mice were stimulated for 2 hours. This short CGRP pulse duration was chosen to capture primary chromatin accessibility changes before downstream transcriptional events could confound the results, whilst preserving acinar cell integrity given their well-established *ex vivo* instability^18^. After stimulation, acinar cells were sorted and subjected to ATAC sequencing. Considering that the increased acinar plasticity observed upon stimulation may reflect a reduced differentiation state, we performed gene set enrichment analysis on ATAC-seq peaks using a curated acinar cell identity signature derived from published transcriptional regulators and functional markers of mature acinar cells (Supplementary table 1)^19–23^. CGRP-treated cells showed reduced enrichment of this signature compared with control acinar cells, which exhibited significant enrichment for this signature (NES=1.98) (Figure 4D). This is consistent with a loss of open chromatin at acinar identity loci that characterizes the mature differentiated state. Amongst the genes that contributed to this phenomenon, we found *PTF1a*, a key acinar cell transcription factor that regulates acinar specification, differentiation and maintenance of identity^19^ (Figure 4E). A similar reduction in open chromatin was found at genes *Bhlha15*, a master co-regulator of acinar cell identity with *Ptf1a*^24^ ,*Rbpjl* and *Spdef* (Figure 4E and S4B and S4C). Collectively, this analysis revealed that CGRP treatment impose a reduction on chromatin accessibility at genes of the core transcriptional machinery responsible for acinar cell commitment, consistent with the maintenance of a more plastic and mitigated terminally differentiated state.

We next performed bulk RNA sequencing on acinar cells following a 2 hour pulse with CGRP and observed a significant early upregulation of biological processes related to development (Figure S4D and S4E). Gene set enrichment analysis revealed a strong enrichment of IL6/JAK/STAT3 signalling, previously associated with epithelial memory of injury^3^, and Epithelial-to-Mesenchymal Transition signature (EMT), linked to ADM and epithelial plasticity^25^ (Figure S4F). Collectively, these changes support that CGRP stimulation limits acinar differentiated state, supporting transcriptional features of injury-induced epithelial reprogramming. Transcriptionally, we observed that the 50 most differentially expressed genes in response to CGRP stimulation were uniformly downregulated, with approximately 50% encoding genes associated with acinar cell function (Figure S4G). In accordance with the ATAC-seq results, amongst the most downregulated genes were *PTF1a* and *Bhlha15*. Supporting a loss of identity commitment, we observed a broad downregulation of genes associated with function and maturation, including those encoding digestive enzymes, zymogen granule proteins and the protein synthesis machinery (Figure 4F).

Collectively, these data suggest that neuronal CGRP stimulation contribute and support pancreatic epithelial memory that is established in the context of inflammation and injury^4^ by directly promoting epithelial plasticity through destabilization of acinar identity and maintaining a reduced cell fate commitment.

### Inhibition of neural memory in males disrupts persistent adaptive tissue memory

Prior pancreatic injury by AP primes the tissue toward an adapted response to subsequent insult. This phenomenon, attributed to incomplete re-establishment of acinar cell identity following the initial injury, enables an accelerated initiation of regenerative programs that can be monitored by the level of ADM^3,4^ (Figure 5A). Our data suggest that this incomplete re-differentiation is maintained by nociceptor signalling in male mice. We therefore asked whether inhibiting nociceptor activity prior to a second AP challenge would restore the epithelial injury response to that of naïve, previously uninjured pancreas.

**Figure 5.**
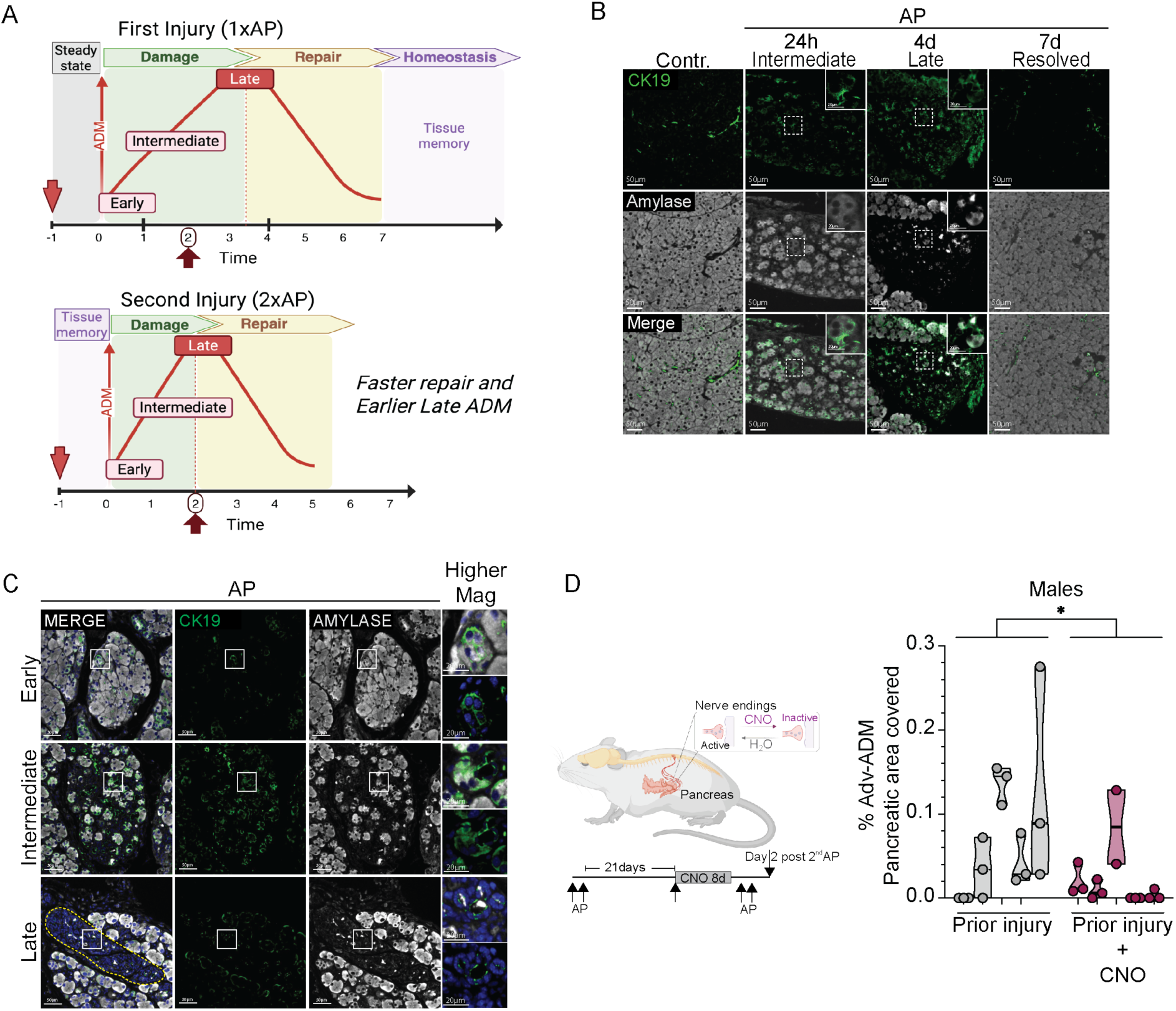
Adaptive tissue memory of injury is reverted by sensory inhibition in males. **(A)** Schematic representation of the kinetics of ADM during the first AP episode (top) and during second AP after repair of the second episode (bottom – re-challenge). Naïve pancreas (top) develops advanced ADM around day 3/4 following the final caerulein injection. Pancreas previously exposed to AP (prior injury), develop a tissue adaptation upon injury repair that accelerates the regenerative response to AP. A second AP episode will anticipate the development of ADM whereby advanced lesions can be found already at 2 days after the final caerulein injection. **(B)** Representative CK19/amylase immunofluorescence images of male pancreas from control and AP animals at 24 hours, 4 days, and 7 days after the final caerulein injection (2-day AP protocol). Scale bars are indicated in the figures. **(C)** Representative CK19/amylase immunofluorescence images at day 2 post injury of male pancreas illustrating early, intermediate, and late ADM stages. Scale bars are indicated in the figures. **(D)** Schematic (left) of transient sensory inhibition followed by AP re-challenge in males. CNO, or vehicle, was delivered in drinking water for 8 days starting 21 days post-AP, prior to a second AP episode. Quantification (right) of parenchyma area covered by advanced/late ADM lesions in animals with (n=5) and without (n=5) sensory inhibition (3 histological sections per animal, 150 μm apart). Data are represented as median (violin plots). Two-way ANOVA. *p < 0.05, Two-way ANOVA See also Figure S5.

In the context of AP, acinar de-differentiation characterizing ADM, a dynamic and heterogeneous process that occurs across a spectrum of cellular states. In naïve animals, as previously described^26,27^, shortly after caerulein administration the ductal marker CK19 is expressed *de novo*, which constitutes the early stages of ADM. At 24 the vast majority of affected areas remain at an early ADM stage, however, in a geographically heterogeneous pattern, ADM progresses to an intermediate stage characterised by loss of intracellular amylase and high CK19 expression. Between 3 and 4 days post-injury the stromal remodelling that accompanies epithelial transdifferentiation expand the interstitial space and disrupt lobular architecture and ADM reaches its advanced stages. The pancreas histologically recovers within one week, with redifferentiation to functional acinar cells and restoration of the tissue is architecture (Figure 5A, top panel, and 5B). In the context of a re-challenge, the dynamics of ADM shift to earlier timepoints, with advanced ADM that can be detected earlier (Figure 5A, bottom panel)^3^.

To interrogate the contribution of nociceptor signalling to this adaptation, we employed Scn10a-Cre;R26-CAG-hM4Di^LSL^ mice, in which the inhibitory DREADD receptor hM4Di is selectively expressed in nociceptors neurons. In these animals, administration of CNO in drinking water drives nociceptor hyperpolarisation and suppression of neurotransmitter release^16^, providing a non-invasive and temporally controlled means of silencing nociceptor activity. This system enabled us to deliver a first AP injury, allow full tissue recovery while selectively silencing nociceptor firing during the post-injury window, and subsequently re-challenge the animals with a second AP insult. Importantly, given the well-established role of sensory innervation in modulating the onset and severity of AP^28^, CNO was withdrawn prior to the re-challenge to release nociceptor inhibition. Due to the dynamic nature of ADM, quantification of ADM as a whole can be confounded by the proportion of acinar cells that initiate but never complete the transition to a ductal phenotype. Therefore, we focused exclusively on areas of advanced ADM, exhibiting complete loss of amylase-positive acinar parenchyma with uniform CK19 expression, indicative of full commitment to a ductal fate (Figure 5C).

Eight days following the first AP injury, animals were placed on CNO-supplemented drinking water for one week to inhibit nociceptor activity. CNO was then withdrawn and animals were re-challenged with a second AP insult (Figure S5A, left panel). Only pancreata of prior-injured animals exhibited a significant presence of advanced ADM coverage following the second insult while, as expected, no significant presence of advanced ADM was detected in pancreas of mice undergoing their first AP episode (naïve) (Figure S5A, right panel). Nociceptor inhibition during the inter-injury recovery window abrogated this adapted response and re-challenged animals did not exhibit a significant presence advanced ADM areas at this early stage (Figure S5A, right panel). This data corroborating a role for nociceptor signalling in the maintenance of tissue injury memory

To confirm that this effect reflected modulation of tissue memory rather than interference with residual tissue repair, we repeated these experiments with nociceptor inhibition initiated three weeks after the first AP injury and directly compared the presence of ADM areas 2 days post injury. Male animals subjected to inter-injury nociceptor inhibition at this later timepoint exhibited a reduced proportion of advanced ADM compared to prior-injured controls (Figure 5D). Notably, no difference between conditions was observed in female animals (Figure S5B) indicating that the regulation of tissue adaptation following injury is different in females mice and that is independently of neural engagement.

Collectively, these data demonstrate that nociceptor signalling, which is selectively engaged in male mice upon injury and maintained as neural-memory, is required for the maintenance of the tissue memory of injury in these animals.

### Transcriptionally distinct neutrophils in female pancreas actively suppress nociceptor activation during injury

Although multiple factors may contribute to sex-specific injury responses and adaptation in the pancreas, we sought here to identify specifically the mechanisms driving the differential engagement of nociceptors during acute pancreatitis in female mice.

Given that AP is characterised by extensive immune cell infiltration and that immune-derived cytokines are established drivers of sensory neuron activation and functional reprogramming^29^, we asked whether the immune compartment of the acutely injured pancreas was sufficient to activate pancreas-innervating nociceptors towards supporting epithelial plasticity. To test this, we generated CM from CD45-positive cells isolated from control adult male pancreas or 24 hours post-AP. Naïve pDRG explants were exposed to this CM for 48 hours, washed thoroughly and subsequently used in co-culture with naïve pancreatic epithelial cells in organoid formation assays. pDRGs primed with AP CM where able to significantly enhanced organoid-forming efficiency compared to control, demonstrating that the immune cytokine milieu of the acutely injured pancreas is sufficient to induce epithelial plasticity boosting activation (Figure 6A).

**Figure 6.**
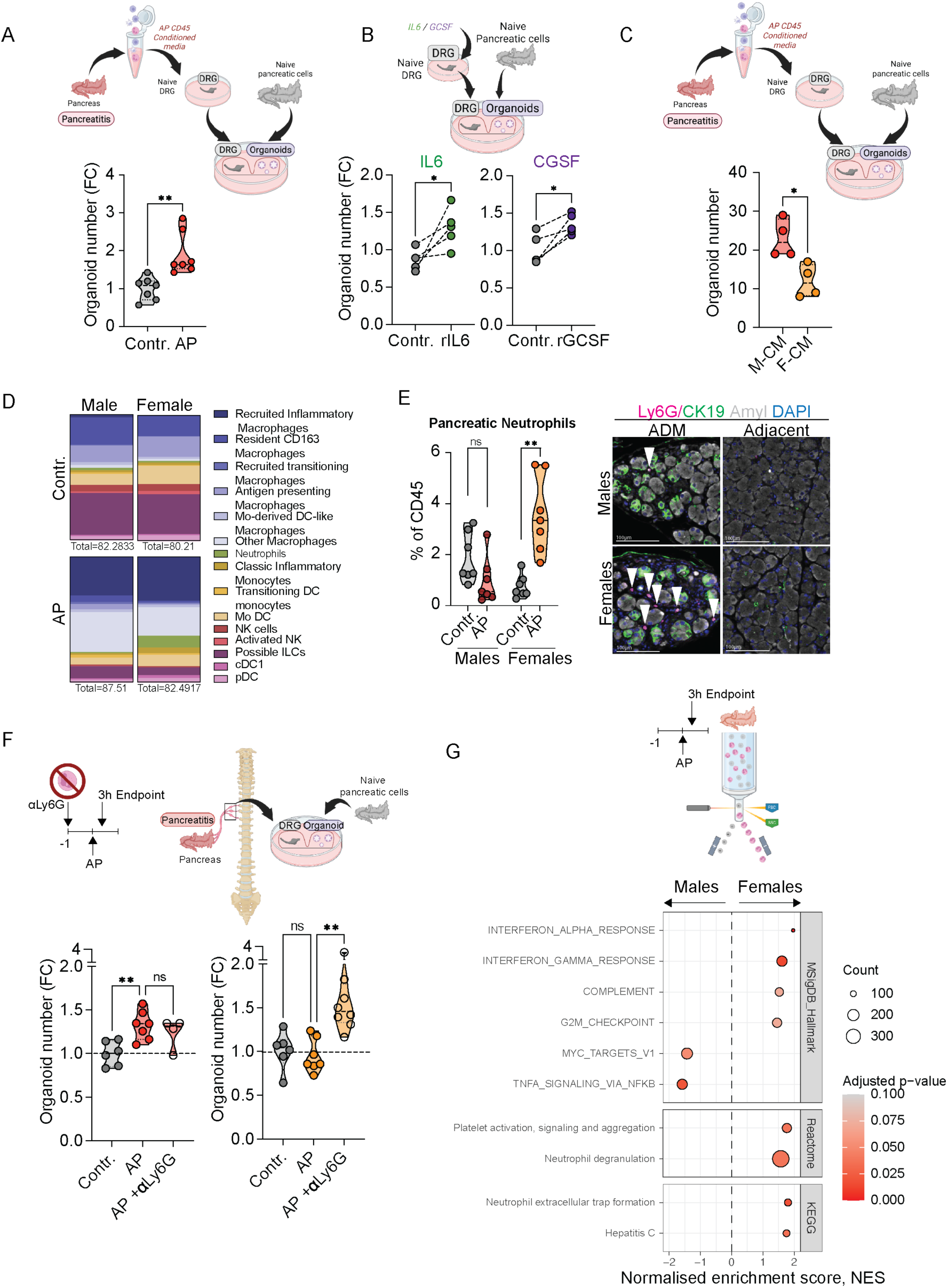
Female neutrophils inhibit sensory neuron activation during AP. **(A)** Schematic (top) of CD45+ pancreatic immune cell secretome assay. CM from CD45+ cells isolated 24 hours post-AP was used to prime pDRGs *ex vivo* before co-culture with naïve epithelial cells in organoid formation assays. Organoid formation efficiency (bottom; control CM n=7, AP CM n=7). Data normalised to control mean per experiment. Data are represented as median (violin plots). **(B)** Schematic (top) of *ex vivo* cytokine treatment of naïve pDRGs. Contralateral pDRGs were treated with vehicle, 200 ng/mL IL-6, or 250 ng/mL G-CSF before co-culture with naïve epithelial cells in organoid formation assays. Organoid formation efficiency (bottom) with control or IL-6-primed pDRGs (left; n=5 per group) and control or G-CSF-primed pDRGs (right; n=5 per group). Data normalised to control mean per experiment. Data are represented as median (violin plots). **(C)** Organoid formation efficiency from epithelial cells co-cultured with pDRGs primed with male (n=4) or female (n=4) AP CD45+ CM, as in (A). Data are represented as median (violin plots). **(D)** Representative FACS profiles of pancreatic-infiltrating immune cells from control and AP male and female mice 24 hours post-injury. **(E)** FACS-based quantification (left) of pancreatic neutrophils (CD45+CD11b+F4/80−Ly6G+) as a percentage of total CD45+ cells in control and AP male and female mice 24 hours post-AP (Male control n=7, Male AP n=7, Female control n=7, Female AP n=7). Data normalised to control mean per experiment. Data are represented as median (violin plots). Unpaired Welch’s t-test. Representative amylase/CK19/Ly6G/DAPI immunofluorescence images (right) from ADM areas of male and female AP mice. Scale bars are indicated in the figures. **(F)** Schematic (left) of neutrophil depletion experiment. Anti-Ly6G (áLy6G) was administered the day before 3 hour AP induction; pDRGs were isolated for co-culture with naïve epithelial cells in organoid formation assays. Organoid formation efficiency (right; Male control n=6, Male AP n=7, Male AP+αLy6G n=4, Female control n=6, Female AP n=7, Female AP+αLy6G n=8). Data normalised to control mean per experiment. Data are represented as median (violin plots).

Previous studies profiling the secretome of CD45-positive cells during the acute phase of pancreatitis identified IL-6 and G-CSF as the main chemokines present^3^. Given that both IL-6 and G-CSF are potent neuromodulators of sensory neuron activity^30,31^, we asked whether these cytokines alone mediate nociceptor priming. Naïve pDRG explants primed *ex vivo* with either recombinant IL-6 or recombinant G-CSF significantly increased the organoid-forming efficiency of naïve pancreatic epithelial cells compared to untreated pDRG (Figure 6B). Supporting a role of G-CSF in pDRG activation, pDRG explants 24 hours post-AP isolated from G-CSF-deficient animals fail to enhance organoid formation from naïve pancreatic epithelial cells, despite equivalent pancreatic tissue damage between those mice (Figure S6A and S6B). These results suggest that IL6 and G-CSF derived from the immune infiltrate of the acutely injured pancreas have an important contribution in driving nociceptor-dependent boosting of epithelial plasticity.

Having identified what drives sensory neuron activation in male animals, we sought to determine what happens in the context of female pancreas. We first compared the capacity of CM from CD45-positive cells isolated from males and females during AP to drive the sensory nerve-dependent boost of epithelial plasticity. We found that pDRG explants primed *ex vivo* with female-derived CD45 CM exhibited a significantly reduced capacity to boost organoids as compared to pDRGs primed with male CM (Figure 6C). This suggested that the lack of nociceptor activation in females could be attributable to a sex-dimorphic immune response. Since IL-6 and G-CSF levels did not differ between sexes in the CD45-positive secretome 24 hours post-AP (Figure S6C), we examined the cellular composition of the immune infiltrate between sexes by flow cytometry. Both sexes exhibited a comparable increase in total immune cell infiltration 24 hours post-AP relative to control pancreas (Figure S6D), however the proportional representation of specific immune cell populations differed between sexes (Figure 6D and S6E). Of these, neutrophils numbers showed the greatest divergence between sexes, with female animals exhibiting a significantly higher neutrophil infiltration 24 hours post-AP (Figure 6E). Notably, within the tissue neutrophils were found in areas of ADM, the same regions where nerve fibre expansion was observed (Figure 6E, right panel), raising the possibility that neutrophils may actively engage with nociceptor activation in females.

To test this, we generated CM from neutrophils isolated from the pancreas of female animals 24 hours post-AP and assessed its effect on IL-6- and G-CSF-mediated nociceptor activation of naïve female pDRG explants. Following 48 hours of stimulation, pDRG explants were washed and co-cultured with naïve pancreatic epithelial cells in organoid-forming conditions (Figure S6F, left panel). Epithelial cells co-cultured with pDRGs activated in the presence of female neutrophil CM failed to boost organoid formation (Figure S6F, right panel), suggesting an inhibitory role of factors derived from female neutrophils on nociceptor activation. To assess this potential inhibition *in vivo*, we examined nociceptor activation in the absence of neutrophils. Neutrophils were depleted in male and female animals one day prior to caerulein administration. To capture neutrophil-mediated regulation of early nociceptor activation, pDRG explants were isolated during injury, 1 hour after the second caerulein injection (3 hour AP protocol, at the start of inflammatory phase) and used in co-culture with naïve pancreatic epithelial cells (Figure 6F, top panel). Strikingly, whilst neutrophil depletion had no effect on the active pDRG-mediated support of epithelial plasticity in males, in females, where no pDRG activation was observed, neutrophil depletion rescued pDRG-mediated support of epithelial plasticity, driving a significant increase in organoid-forming efficiency (Figure 6F). These data support the notion that neutrophils, specifically in female pancreas, actively suppress sensory activation upon AP. Bulk RNA sequencing of neutrophils isolated from pancreata of male and female animals at the start of inflammation confirmed a transcriptionally distinct profile between sexes (Figure 6G), suggesting that the female-specific inhibitory capacity is underpinned by a sex-dimorphic neutrophil transcriptional program.

Collectively, these findings demonstrate that during AP, IL-6 and G-CSF within the inflammatory milieu trigger a neuronal activation and consequent neural-memory, which supports the long-term epithelial plasticity characteristic of tissue memory. In females, however, a transcriptionally distinct neutrophil population actively suppresses this response, positioning neutrophils as sex-specific gatekeepers of neural memory of injury.

## DISCUSSION

We investigated the tissue features underlying the persistent memory of pancreatic injury that enhances resilience to secondary insults. Acute pancreatitis (AP) triggers a rapid dedifferentiation of acinar cells accompanied by the reactivation of regenerative and progenitor-like programs^32^. Previous studies have shown that these regenerative states can persist long after histological repair and resolution of inflammation, establishing a metastable epithelial state with enhanced regenerative potential^3,4^. Because inflammatory epithelial memory relies on the sustained maintenance of progenitor-like activity, it is likely that, at the tissue level, this state is regulated by signals from the surrounding niche. We therefore explored whether additional cellular compartments retain a persistent memory of injury that could function together with epithelial memory to sustain long-term tissue adaptation and resilience.

Here, we investigated the pancreatic response to AP in mature 6-month-old male and female mice. While no major sex-dependent differences were observed in tissue injury or histological repair, male mice displayed increased epithelial plasticity after tissue recovery, as measured by enhanced *ex vivo* organoid-forming capacity (Figure 1F,G). This phenotype correlated with the presence of pERK-positive cells within the intra-acinar space that were closely associated with neural fibres (Figure 1H).

During injury, both male and female pancreata exhibited neural remodelling characterised by increased neural density, particularly within regions undergoing acinar-to-ductal metaplasia (ADM). However, increased CGRP-positive fibres were detected preferentially in male pancreata, accompanied by a concomitant increase in CGRP expression in pancreas-innervating dorsal root ganglia (pDRGs) (Figure 2).

The peripheral nervous system (PNS) has increasingly been recognised as a key determinant of pancreatic disease progression and outcome^11^. However, its involvement in AP is still poorly understood. A growing body of literature has positioned CGRP as an emerging regulator of epithelial and cancer cell biology^33–36^, however, its direct role in pancreatic epithelial plasticity has not previously been investigated. Here, we demonstrate that activation of pDRGs following tissue injury enhances epithelial plasticity *ex vivo* in organoid co-culture systems and direct chemogenetic-activation of sensory fibres *in vivo* increased epithelial plasticity independently of the injury. Together, these findings establish a direct functional link between sensory neuronal activity and pancreatic epithelial plasticity (Figure 3A-C). Most importantly, the neural activity associated with enhanced epithelial plasticity persisted after tissue repair as neural-memory of the injury, identifying sensory neurons as an additional tissue niche compartment capable of retaining a lasting memory state. Furthermore, sensory ablation following tissue repair reversed the epithelial memory of injury, as evidenced by a reduction in epithelial plasticity (Figure 3D-H). Although we did not directly assess the contribution of CGRP signalling in the context of acute pancreatitis, direct stimulation of naïve epithelial cells with CGRP was sufficient to induce epigenetic and transcriptional changes (Figure 4D–F), suggesting that neural-memory via CGRP may preserve epithelial plasticity long term by restraining acinar lineage commitment. Together, these findings define the existence of a long-term neural memory that is functionally coupled to epithelial tissue memory.

Interestingly, female mice do not display elevated CGRP^+^ sensory fibres, and, consistently, no neural memory was observed in mature female pancreas. Here, while the steady state tissue homeostasis and epithelial regenerative activity between male and female was not investigated, we found that the level of epithelial plasticity after tissue repair in female was reduced compared to male when measured *ex vivo* using organoid formation assay (Figure 1F). However, females pancreata retained a functional tissue memory of the injury which was independent of neural memory (Figure S5B), suggesting that other mechanisms of tissue-mediated adaptation to the injury might engage in this female pancreas.

Sex differences in the transcriptional profile of DRG sensory neurons in response to peripheral nerve injury have been previously reported^37^, and sex-dependent differences in the immune responses are well known^38^. When analysing the immune infiltrate in male and female pancreas during AP, we observed differences in the immune cellular profile with a higher presence of the myeloid compartment in female pancreas, particularly neutrophils. Those neutrophils showed a different activation signature and were functionally able to inhibit pDRG activation *in vitro* and *in vivo* (Figure 6). Strikingly, in patients with pancreatitis, sensory stimulation was found to be reduced in females compared with males, as assessed by skin sensitivity within the dermatomal segments innervated by the same DRGs that project to the pancreas^39^. These findings suggest the existence of an active suppression of sensory signalling in female patients in the context of pancreatic injury. Our data could therefore suggest that specific neutrophil responses to pancreatitis specifically in females might be responsible for this reduced sensory activation.

Together, our findings identify a previously unrecognised neuro-epithelial axis that sustains tissue memory long-term following pancreatic injury and suggests that this neural memory is an important component of the epithelial tissue niche that provide active support for regenerative competence. We also propose that sex-specific immune responses shape the persistence of sensory neuronal activation, thereby determining the extent of epithelial memory and regenerative plasticity maintained after tissue repair. Given that increased epithelial plasticity may create a tissue state more permissive to cancer predisposition^3,4^, this sex-dimorphic effect, marked by a more pronounced epithelial plasticity in males compared with females, could be functionally linked to the higher incidence of pancreatic cancer in men^40^. These results position neural memory as a key component of long-term tissue adaptation and reveal the tissue microenvironment as an active regulator of epithelial resilience after injury.

## MATERIALS AND METHODS – STAR METHODS

### EXPERIMENTAL MODEL AND STUDY PARTICIPANT DETAILS

#### Mouse models

All mice were bred and maintained under specific-pathogen-free conditions and provided with food and water *ad libitum* by The Francis Crick Biological Research Facility.

Wild-type C57BL/6Jax from The Jackson Laboratory were provided by The Francis Crick Biological Research Facility.

Scn10a-Cre; R26^iDTR^ animals were maintained on a C57BL/6 background and obtained by crossing homozygous Rosa26-LSL-iDTR with homozygous Scn10a-Cre mice. Rosa26-LSL-iDTR were provided by The Francis Crick Biological Research Facility. These animals harbour a loxP-flanked STOP cassette upstream of the open reading frame of the simian Diphtheria Toxin Receptor (DTR). Scn10a-Cre animals were kindly provided by Leanne Li from the Francis Crick Institute. These animals harbour a Cre recombinase sequence at the 3’ UTR of Scn10a. Cre expression starts at E14 and is restricted to small-diameter neurons in dorsal root, trigeminal and nodose ganglia, specifically in Nav1.8-expressing sensory nociceptor neurons(Nassar 2004) . Sensory neurons remain intact and functional until administration of Diphtheria Toxin, from which point nociceptor neurons are irreversibly ablated.

Scn10a-Cre animals (Strain #:036564, The Jackson Laboratory) were used for intrathecal administration of virus. These animals were maintained on a C57BL/6 background.

R26-CAG-hM4Di^LSL^; Scn10a-Cre animals were maintained on a C57BL/6 background and obtained by crossing heterozygous R26-CAG-hM4Di^LSL^ with homozygous Scn10a-Cre mice. R26-CAG-hM4Di^LSL^ mice harbour a human M4 muscarin 4 receptor sequenced modified to abolish receptor affinity for the native ligand, acetylcholine (ACh), but allow receptor binding and subsequent activation by the small drug-like molecule clozapine-N-oxide (CNO) (Zhu 2016) . In these animals, CNO administration hyperpolarizes nociceptor sensory neurons, decreasing excitability and suppressing neurotransmitter release at the presynaptic terminals.

GCSF KO animals maintained homozygous on a FVB background. These animals lack exons 1-3 of the GCSF gene replaced by a lacZ gene and neomycin resistance cassette rendering deficient for the GCSF protein (Lieschke 1994).

Both male and female mice were used in this study. Specification of sex is included in the figure legends of the respective experiment. Unless stated otherwise, animals were used between 16 and 22 weeks of age. All experiments were performed controlling for age, sex and litter. Breeding and all animal procedures were performed at the Francis Crick Institute in accordance with UK Home Office regulations under project license PP5920580.

#### Dorsal root ganglia explants

Dorsal root ganglia (DRG) were isolated from adult mice following a previously described dissection protocol (Sleigh 2016,Sleigh 2020). Briefly, following euthanasia, the spinal column was excised and cleared of surrounding musculature, fat and soft tissue. The column was cut longitudinally along the midline and pinned medial side up in Sylguard184 silicone-lined petri dishes flooded with ice-cold -Ca,-Mg HBSS (Fisher Scientific). From the Sleigh et al. 2016 protocol that we already have:

Under a stereoscope, the spinal cord and meninges were carefully removed to expose the DRG, which were extracted by grasping the distally projecting axon bundles with fine forceps. Residual meninges and axon bundles were trimmed using fine spring scissors. Thoracic DRG at level T10 and lumbar DRG at level L1 were identified using the most caudal rib pair as an anatomical landmark (T13), with T10 located three levels rostral to the last rib and L1 identified as the first lumbar ganglion caudal to the thoracic column. Isolated ganglia were maintained as intact explants in ice-cold -Ca,-Mg HBSS until further processing. DRG explants were subsequently used for ex vivo priming, co-culture experiments with primary epithelial cells or fixed-frozen immunofluorescence staining.

Sylguard 184 silicone elastomer (Dow Corning) was prepared by combining the elastomer base with the curing agent at a 10:1 ratio. The mixture was used to line petri dishes and allowed to set for a minimum of 48 hours at room temperature. To remove bubbles prior to setting, a vacuum desiccator was applied.

#### Primary pancreatic epithelial organoids

To generate primary pancreatic epithelial organoids the pancreatic tail and body of the pancreas was isolated using fine forceps and scissors and collected in ice cold -Ca,-Mg HBSS. To minimise mechanical damage to the pancreatic parenchyma, the body and tail of the pancreas were isolated by indirect manipulation, grasping the adjacent spleen with fine forceps rather than the pancreas itself. Pancreatic tissue was finely minced using two crossed scalpel blades into small fragments and enzymatically digested in 20mL of digestion medium composed of Advanced DMEM/F12 and 1mg/mL collagenase type V from Clostridium histolyticum (Sigma-Merck). Digestion was performed for 20 minutes in a 37°C shaker at 140rpm. The reaction was stopped by adding 30mL of ice-cold MACS buffer containing (0.5% BSA (Sigma) and 250 mM EDTA (Sigma) in PBS). To obtain a single cell suspension, samples were centrifuged for 4 min at 400xg at 4°C, and the pellet resuspended in MACS buffer and filtered twice with a 70 um and 40μm filters sequentially.

For assessment of pancreatic intrinsic plasticity, freshly dissociated acinar cells were manually counted using a hemocytometer and 10×10^4^ cells were seeded in a 25 μl drop of growth factor reduced (GFR) Matrigel (Corning) in a 24-well cell culture plate. The combined cell-matrigel drop was allowed to set at 37 °C for 30 minutes before being supplemented and maintained in a medium of Advanced DMEM containing: pencillin/streptomycin (1:100), 50 ng/ml EGF (Peprotech), 100 ng/ml FGF10 (Peprotech), 10 nM Gastrin I (Sigma), 10% Noggin-CM (Cell Sciences, Francis Crick Institute), 10% RSPO-CM (Cell Sciences, Francis Crick Institute), 1.25 mM N-acetylcysteine (Sigma), 10 mM Nicotimanide (Sigma), 1X B27 Supplement (Fisher Scientific), as adapted from (Huch 2013). On the day of seeding, cultures were supplemented with 10.5 μM RhoK inhibitor Y-27632 (Stratech) to help cells cope with stress. Media were changed every 2-3 days, from which moment Y-27632 was no longer added. The number of organoids in each well were recorded 10 days post seeding using an EVOS microscope (Thermo Fisher Scientific) and quantified in Fiji.

#### Dorsal root ganglia monocultures and co-cultures with pancreatic epithelial cells/organoids

For assessment of DRG-dependent plasticity, DRG explants isolated as described above, were placed on top of a 5 μL drop of GFR Matrigel (Corning) on one side of the well of a 24-well cell culture plate. The DRG-containing Matrigel drop was allowed to set for 15 minutes at 37 °C before 10×10^4^ cells were seeded in a 15 μl drop of growth factor reduced (GFR) Matrigel (Corning) in the opposite side of the well, ensuring physical separation between DRGs and epithelial cells. Cultures were maintained in organoid media, described above, without growth factors EGF, FGF and Gastrin (referred to DRG media) and supplemented with 0.5 μM RhoK inhibitor Y-27632 (Stratech) for the first 3 days of culture, after which cultures were transitioned to maintained in complete organoid media without RhoK and changed every 3 days. The number of organoids in each well were recorded 10 days post seeding using an EVOS microscope (Thermo Fisher Scientific) and quantified in Fiji.

For DRG to ex treatments, DRG explants, isolated as described above, were kept in monocultures, placed on top of a 5 μL drop of GFR Matrigel (Corning) on a 96-well plate. The DRG-containing Matrigel drop was allowed to set for 15 minutes at 37 °C and DRG media, containing treatment agents (see DRG ex vivo treatment), was added.

### METHOD DETAILS

#### Acute pancreatitis

Acute pancreatitis was induced by injecting 50 μg kg^-1^ of caerulein (Sigma), dissolved in endotoxin free PBS into the peritoneum of animals, every hour for 6 hours, during a period of 2 days (12 injections in total). Unless stated otherwise in the figure legends, all animals were between 16 and 22 weeks of age. Sex specified in figure legend. Acute pancreatitis was induced in three different settings: full protocol on treatment-naive animals (AP); full protocol of animals previously subjected to a full acute pancreatitis protocol (AP re-challenge experiments); induction of early phase acute pancreatitis (3-hour AP) for assessment of neutrophil contribution to neuronal support of epithelial plasticity and neutrophil RNA sequencing.

Re-challenge of acute pancreatitis protocol was performed as the first event after recovery of the first acute protocol. Induction of early injury phase was achieved by 2 injections of 50 μg kg^-1^ of caerulein (Sigma), dissolved in endotoxin free PBS into the peritoneum of animals, one hour apart of each other. Experiment was terminated one hour after the last injection rendering the protocol a 3-hour AP.

#### Tissue fixation and storage

Mouse tissue samples were fixed overnight for pancreas, 8h for DRGs, in 10% neutral-buffered formalin (NBF) and transferred 70% ethanol until further processing. Samples were either embedded in paraffin block for long term storage at room temperature or cryoprotected in a sucrose gradient OCT embedding. For the latter, after fixation, samples were transferred to 10% sucrose (Sigma) in PBS, followed by 20% and 30%, with each incubation lasting no less than 16 hours. Samples in 30% sucrose were embedded in OCT (CellPath). OCT-embedded tissues were placed in cryomolds, oriented as required, and snap-frozen by floating the mold on a dry ice/ethanol bath until the OCT turned uniformly white and opaque, after which blocks were stored at −80°C until sectioning. Paraffin and frozen sections were cut using a microtome (Leica) and mounted onto Superfrost Plus slides.

#### Immunofluorescence on paraffin samples

Tissue sections of 4 μm thickness were cut using a Leica microtome and mounted onto Superfrost Plus slides. For each animal, 3 levels were cut and stained 150 μm apart. Slides were incubated for 1 hour at 60°C to increase adherence and then deparaffinized and rehydrated using standard methods. Heat mediated antigen retrieval was achieved by pre-heating 10 mM pH6 tri-Sodium citrate dihydrate (Sigma) solution for 8 minutes full power in a commercial microwave oven (Maestrowave MW10) and then incubation tissue slides in the pre-heated solution for 15 minutes in the microwave at medium power. Slide were allowed to cool down under running tap water until reaching room temperature and blocked for 45 minutes with blocking solution composed of 1% BSA (Sigma), 10% FBS (Fisher Scientific), 0.2% Triton X-100 (Sigma). For stains using mouse monoclonal antbodies, blocking buffer additionally contained FcR Blocking reagent 1:100 (Miltenyi). Primary antibody incubation was performed overnight in blocking serum at 4°C inside a humid chamber. The following day, slides were washed three time for 5 minutes in 1X PBS, incubated with fluorescently labelled secondary antibodies and 1 ug/mL DAPI in blocking serum at room temperature for two hours and mounted after three additional washed with 1X PBS. Slides were scanned with Akoya Phenolmager slide scanner and analysed in QuPath-0.6.0.-arm64.

The following antibodies and dilutions were used: Amylase (goat, #sc-12821, Santa cruz, 1:300); Amylase (mouse, #sc-46657, Santa Cruz, 1:300); STMN1 (rabbit, clone EP1573Y, #ab52630, 1:500); Phospho-p44/42 MAPK (Erk1/2) (Thr202/Tyr204) (rabbit, #4370, Cell signaling, 1:300), Troma-III (CK19) (rat, MABT913, Merck, 1:300), βIII Tubulin (rabbit, EP1569Y, ab52623, abcam, 1:250); βIII Tubulin (TUJ-1, mouse, ab14545, abcam, 1:200); αSMA (mouse, #A2547, Merk, 1:200); Ki67 (rabbit, ab15580, abcam, 1:300); phosphorylated histone 3 - pH3 (rabbit, #9701, cell signaling, 1:300). All secondary antibodies were raised in donkey (Invitrogen) and used at a concentration of 1:200 in blocking serum against the species of the primary antibody.

Triple immunofluorescence for Ly6G, CK19 and Amylase as performed on pre-baked 4 μm paraffin tissue sections of thickness using a Leica Bond Rx automated stainer. The following antibodies and dilutions were used: amylase (Bios Antibodies, # BS-4030R, 1:100); CK19 (Development Studies Hybridoma Bank, #TROMA-III-c, 1:100); Ly6G (Cell Signalling Technology, #87048S, 1:100). Antigen retrieval was performed using AR9640, (Leica) for 20 min at 95°C before each staining cycle. Detection was performed using Novolink Polymer Detection Systems (Leica , # RE7260-CE) with opal 570 (FP1488001KT, 1:500), Opal 690 (FP1497001KT, 1:150) and Opal 520 (FP1487001KT, 1:500). Slides were counterstained with DAPI (Thermo Scientific, #62248, 1:2500) and mounted using ProLong™ Gold Antifade Mountant (Thermo Fisher, P36934). Slides were scanned with Akoya Phenolmager slide scanner and analysed in QuPath-0.6.0.-arm64 Analysis details in Quantification and Statistical analysis.

#### Haematoxylin and Eosin staining

Tissue sections of 4 μm thickness were cut using a Leica microtome and mounted onto Superfrost Plus slides. Slides were incubated for 1 hour at 60°C to increase adherence and then deparaffinized and rehydrated using standard methods. Haematoxylin and Eosin stain, as well as coversliping, was performed using a Tissue-Tek Prisma Plus automated stainer.

#### Immmunohistochemistry

Tissue sections of 4 μm thickness were cut using a Leica microtome, mounted onto Superfrost Plus slides and baked for 1hr at 60°C before staining. Heat mediated antigen retrieval was performed using pH9 Tris-EDTA buffer for 18 minutes in the microwave on 80% power. After cooling slides in slow running water they were incubated in 1.6% H202/PBS for 12 minutes to quench endogenous peroxidase activity. Samples were delineated using hydrophobic barrier pen prior to blocking in 1% BSA/2.5% Horse Serum/PBS for 1 hour at room temperature. Primary antibody CD45 (Cell Signalling, 70257S, Clone D3F8Q 1:250) was then incubated overnight at 4 °C prior to washing with PBS. Secondary antibody (ImmPress Horse anti-Rabbit HRP-Polymer, MP-7401-15) was added and incubated at room temperature for 45 minutes prior to washing with PBS. DAB substrate (Vector Labs, SK4100) was prepared using manufacturer’s instructions and incubated for 4 minutes before washing with water. Slides were counterstained with Haematoxylin and coverslipped using TissueTek Prisma plus automated stainer.

Stained slides were scanned using Zeiss AxioScan.Z1 and quantified using QuPath-0.7.0.-x64.

Analysis details in Quantification and Statistical analysis.

#### Immunofluorescence on frozen tissue frozen tissue

For DRG immunofluorescence, frozen tissue blocks were sectioned using a cryostat (Leica) maintained at a cabinet temperature of −20°C and an object temperature of −18°C. Sections of 4 μm thickness were cut and mounted onto SuperFrost Plus slides, allowed to air dry at room temperature for 30–60 minutes and stored at −80°C until further processing. Stored sections were placed at room temperature for 30 minutes, followed by a 30-minute incubation with 1X PBS and a 1-hour incubation with blocking serum composed of 1% BSA (Sigma), 10% FBS (Fisher Scientific), 0.2% Triton X-100 (Sigma). Primary antibodies were incubated overnight in blocking solution, washed three times for 5 minutes in 1X PBS and incubated with secondary antibodies (1:200) and 1 μg/mL DAPI in blocking serum at room temperature for two hours. Following a set of three washes for 5 minutes with 1X PBS, slides were mounted, allowed to dray for a night and scanned with Akoya Phenolmager slide scanner. Analysis was performed in QuPath-0.6.0.-arm64. Primary antibodies used: CGRP (goat, ab36001, abcam, 1:200) and βIII Tubulin (rabbit, EP1569Y, ab52623, abcam, 1:250). Secondary antibodies were raised in donkey (Invitrogen) and used at a concentration of 1:200 in blocking serum against the species of the primary antibody.

For CGRP fibre quantification on cryoprotected pancreatic frozen samples, OCT blocks were sectioned using a cryostat (Leica) maintained at a cabinet temperature of −20°C and an object temperature of −18°C. Sections of 100 μm thickness were cut and mounted onto SuperFrost Plus slides, allowed to air dry at room temperature for 30–60 minutes and stored at −80°C until further processing. Prior to staining, slides were washed with 1X PBS for 30 minutes at room temperature and the tissue was incubated with a 0.24% NH4Cl (Sigma) PBS quenching solution for 10 minutes. Tissue was permeabilized and saturated for two hours at room temperature in blocking solution containing 15% donkey serum (Sigma), 0.20% glycine (Sigma), 2% BSA (Sigma), 0.25% pork skin gelatine (Sigma), 0.5% Triton X-100 (Sigma) in PBS. Primary antibodies were incubated for 24-hours in blocking solution overnight at 4°C (CGRP (anti-goat Abcam, #ab36001, 1:200), anti-β3-Tubulin (anti-rabbit Abcam, #ab52623, 1:250)) in a humid chamber Following primary antibody incubation, slides were washed there times in 1X PBS at room temperature and an additional 24-hour wash at 4°C in a humid chamber. Fluorescently-labelled secondary antibody incubation was performed for two hours at room temperature (488 donkey anti-rabbit for B3Tub 1:250 and 555 donkey anti-goat for CGRP 1:200 diluted in PBS) followed by a 5 minute incubation with DAPI (Thermo Fisher) at 1μg/mL in PBS. Slides were washed as before and mounted using Dako fluorescence mounting medium (Agilent).

Analysis details in Quantification and Statistical analysis.

***Fast light-microscopic analysis of antibody-stained whole organs (FLASH)*** Whole pancreas imaging was performed by Fast light-microscopic analysis of antibody-stained whole organs (FLASH) as described in (Messal 2021). Animals were culled and cardiac perfusion of 20mL of 1X PBS was performed by making a small incision in the right atrium and perfuse with 20 ml of PBS through the left ventricle. A second perfusion with 20 ml of 4% (wt/vol) PFA (Thermo Fischer) and the pancreas was removed with fine surgical tools avoiding damage to the tissue. The isolated organ was fixed overnight at room temperature in 50mL of 4% (wt/vol) PFA (Thermo Fischer), washed for 10 minutes in 50 mL of PBS at room temperature and incubated first for 1-hour at room temperature and then for 12h at 54°C in an SDS-based antigen retrieval solution prepared by addinf 4% SDS (Sigma) to 200mM Borate (Sigma). Following antigen retrieval, samples were washed three times for 1 hours in 50mL of 1X PBS and moved to a Wheaton vial for subsequent steps. Blocking was performed for 1 hour at room temperature in a solution containing 10% FBS (Fisher Scientific), 0.02% sodium azide (Sigma), 1% BSA (Sigma) and 5% DMSO (Sigma). Primary antibodies were added to the solution (Troma-III (CK19) (rat, MABT913, Merck, 1:100), and βIII Tubulin (rabbit, EP1569Y, ab52623, abcam, 1:250) and incubated for two consecutive nights at room temperature. Samples were then washed three times for 20 minutes in 9mL of 1X PBS (Sigma) and incubated for two additional nights with fluorescently-labelled secondary antibodies (donkey anti-rat 488 and donkey anti-rabbit 555 - Invitrogen) at room temperature in the dark. Following another three 20-minute washed in 1X PBS, dehydration and tissue clearing was performed by transferring the samples to watch glasses in 100% (vol/vol) methanol (Sigma) and gradually increasing concentration of methyl salicylate (Merk) by progressive 30-minute incubations with of 25%, 50%, 75% and 100% and 100% methyl salicylate (Merk) in methanol. Cleared and stained samples were imaged by light-sheet LaVision light-sheet and rendered with respective software.

All incubation steps were performed on a nutator.

#### rAAV production

Transfer plasmids pAAV CAGs-DIO hM3Dq-mCherry (expressing cre-dependent activatory DREADD and mCherry) and pAAV EF1a Flex mCherry (expressing cre-dependent mCherry control) were obtained from the Francis Crick Institute’s Viral Vector core facility and packaged into adeno-associated virus (AAV) serotype PHP.S particles by triple transduction of HEK-293T cells as described previously (Challis 2019), with minor modifications. HEK-293T cells, maintained in DMEM + 10% FBS (Fisher Scientific) + Pen/Strep, were passaged the day before transfection and seeded at 6-7·10^6^ cells per 10-cm dish. Cells were transfected with pHelper : pUCmini-iCAP-PHP.S : transfer plasmid in a 1:1.3:1 molar ratio using PEI-MAX (24765, Polysciences Inc.). pHelper was from Agilent Technology; pUCmini-iCAP-PHP.S was a gift from Viviana Gradinaru (Addgene plasmid #103006) (Chan 2017). After 6 hours media was changed. Supernatant collected 3 days after the transfection was pooled with supernatant and cells collected at day 5 after transfection; cells were lysed by adding 2.5 ml chloroform (288306, Merck), vortexed for 2 min, then centrifuged at 3000 g for 20 min. Supernatants were precipitated with 8% PEG8000 (81268, Merck) + 82mM NaCl overnight at 4°C. AAV particles were pelleted by centrifugation at 3000 g for 20 min. The pellets were resuspended in 4 ml gradient buffer (10 mM Tris pH 7.6, 150 mM NaCl, 10 mM MgCl_2_) containing 50 U/ml Denarase (20804, c-Lecta) and incubated 37°C for 45 min; 2.5 ml chloroform were added, followed by vortexing for 2 min and centrifugation at 3000 g for 20 min. The aqueous layer was collected and loaded onto a Iodixanol (07820, StemCell Technologies) step gradient, centrifuged for 2 hr at 52000 RPM in a Type 70.1 rotor. The 40% layer was collected and filtered through a 0.45 µm syringe filter onto a 4 ml Amicon-100 filter (UFC810024, Millipore), pre-wetted with 0.01% Pluronic Acid (24040, Invitrogen) at RT for 1h, to concentrate and buffer exchange with storage buffer (1x PBS; 5% Sorbitol, 0.1M NaCl [0.25M NaCl final]).

AAVs were titrated using the method described in (Aurnhammer 2012), on a Quant3 (Perkin Elmer) using PowerUp SYBRgreen master mix (A25742, Perkin Elmer) and a linearized plasmid used as a standard curve.

#### Intrathecal injection of rAAV and chemogenetic stimulation

Scn10a-cre males aged between 11- to 15-week-old received intrathecal AAV administration between lumbar vertebrae L4–L6 under isoflurane anaesthesia. Mice were administered AAV CAGs-DIO hM3Dq-mCherry or control AAV at a dose of 2·1010 viral genomes per injection in a volume of 10 μl, with two injections delivered 3 days apart. Animals were monitored for at least 4 weeks following AAV administration to allow transgene expression in Scn10a+ DRG sensory neurons. Mice subsequently received intraperitoneal injections of 1 mg/kg Clozapine N-oxide hydrochloride (CNO, 5780-25mg-CAY, Cayman Chemical), once daily for 3 consecutive days. One day after the last CNO administration, pancreatic epithelial cells were isolated and tested for organoid formation efficiency as described above.

#### Genetic sensory nerve depletion

Genetic nociceptor nerve depletion was achieved employing the Scn10a-Cre; R26^iDTR^ model. In these animals, nociceptor neurons express the simian Diphtheria Toxin Receptor (DTR) which enables conditional and irreversible ablation of nociceptor neurons upon treatment with diphtheria toxin (DT). Nerve deletion was performed in there different experimental settings: 3 days, 7 days and 21 days after the last caerulein injection (acute pancreatitis protocol described above). In all three experimental conditions, a working solution of 2 μg/mL of DT (Sigma) in 1X PBS was prepared and 5 μl per gram of mouse body weigh was injected intraperitoneally (10 μg/kg) once a day for 3 consecutive days. Animals were kept no longer that 11 days after that. Same dose was used in WT mice with no measurable adverse effects.

#### DRG conditioned medium on epithelial cell plasticity

Acute pancreatitis was induced in C57BL/6 16-22-week-old male mice as described. Twenty-four hours after the last caerulein injection, pancreatic innervating T10 DRGs (2 per animal) of AP and control mice were removed, as described above, placed in a 5 μl drop of GFR Matrigel (Corning) on a 28-well plate until the Matrigel was set and 200 μl of DRG media (described in DRG monocultures) were added. Following 3 days of culture, media was collected, centrifuged once at 300g for 10 minutes at 4°C, and the supernatant was subsequently centrifuged for 10 minutes at 2000g at 4°C and re-collected. Conditioned medium was mixed 1:1 with fresh DRG media and used to culture 10×10^4^ freshy isolated pancreatic epithelial cells on a 25uL GFR Matrigel drop in a 28-well plate. Media was supplemented with 0.5 μM RhoK inhibitor Y-27632 (Stratech) and was not changed until the end of the experiment.

#### Immune cell conditioned medium generation

Conditioned medium was generated from CD45-positive cells and neutrophils using the same general protocol, with cell-type specific modifications described below. Pancreas was collected from experimental animals, finely minced into small fragments and enzymatically digested in 20mL of digestion medium composed of Advanced DMEM/F12 and 1mg/mL collagenase type V from Clostridium histolyticum (Sigma-Merck). Digestion was performed for 20 minutes in a 37°C shaker at 140rpm and stopped by adding 30mL of ice-cold MACS buffer containing (0.5% BSA (Sigma) and 250 mM EDTA (Sigma) in PBS). Single cell suspension was centrifuged for 4 min at 400xg at 4°C, the pellet resuspended in MACS buffer and filtered twice with a 70 um and 40μm filters sequentially. Pancreatic cell digestion leads to excessive acinar cell death which reduces efficiency of downstream steps. To remove these from the suspension, the EasySep Dead Cell Removal Kit (Annexin V, Miltenyi) was used according to manufacturer instructions. Briefly, cells were counted and prepared to a concentration of 1×10^8^ cells per mL of MACS buffer for a maximum volume of 2mL. per mL of sample, 50 μl of Dead Cell Removal Cocktail was added, followed by 50

μl/mL of sample of Biotin Selection Cocktail. Samples were mixed and incubated for 3 minutes at room temperature. A volume of 100 μl/ml of sample of, previously vortexed, RapidSpheres^TM^ was added and immediately supplemented with MACS buffer to a maximum total volume of 2.5 mL. Tubes were placed in an EasySep magnet and separation was allowed to occur for 3 minutes. Following incubation, the enriched suspension was collected and centrifuged at 300g for 10 minutes at 4°C. Cell pellet was resuspended in 100μl of MACS buffer with 1:50 FcR blocking reagent (Miltenyi) and incubated for 10 minutes on ice. Following centrifugation at 300g for 5 minutes, cells were resuspended in 300 μl of MACS buffer and 30 μl of CD45 Microbreads (Miltenyi) (for CD45 Conditioned medium) or Ly6G microbeads UltraPure (for neutrophil conditioned medium) and incubated on ice for 20 minutes. Following bead incubation, samples were washed with 2 mL of MACS buffer, centrifuged at 300g for 10 minutes at at 4°C and resuspended in 500 μl of MACS for positive selection magnetic cell sorting (MACS, Miltenyi) using LS columns (Miltenyi) followed by MS column (Milteny) separation according to manufactors instructions. For LS separation, columns were placed on a Miltenyi MACS Separators and equilibrated with 3 mL of MACS, allowing the solution to pass by gravity. Cell suspention was added and washed three times with 3 mL MACS, allowing the column to empty by gravity each time. After the third was, the column was removed from the magnet stand onto a falcon, 5 mL of MACS were added to the columns and cells were released by plunging in one single push. Suspention was then MACS positively with MS columns selected to guarantee purity. Protocol was as for LS with reduced volumes of 1mL. Once positive selection was complete, cells were counted with Countess II Automated Cell Counter (Thermo Fisher) and 1×10^6^ cells were plated in AdDMEM/F12 (Thermo fisher) supplemented with 1:250 10X B27 (Thermo Fisher). No longer than 16 hours later, media was collected, centrifuged once at 300g, followed by a centrifugation at 2000g, both for 10 minutes at 4°C, and the media either used fresh for priming of DRGs or snap frozen for ELISAs.

#### CGRP stimulation protocols

CGRP stimulation was performed with a synthetic rat/mouse α-CGRP peptide (Biosynth, #PCG-4163-V; CAS 83651-90-5) at a concentration of 10nM in water. The timing of stimulation and downstream steps following incubation varied according to the aim of the experiment.

To assess CGRP effect on functional pancreatic plasticity, pancreatic epithelial cells were isolated as performed for organoid cultures until obtaining a single cell suspension and 1×10^4^ cells epithelial cells were seeded in a 25 μl drop of GFR Matrigel (Corning) in a 24-well cell culture plate. The combined cell-matrigel drop was allowed to set at 37 °C for 30 minutes and organoid media (described above) with 10nM CGRP or vehicle H_2_O control was added. On the day of seeding, cultures were supplemented with 10.5 μM RhoK inhibitor Y-27632 (Stratech). Media were changed every 2 days containing CGRP or H_2_O vehicle. From day 2 of culture, Y-27632 was no longer added. The number of organoids in each well were quantified 10 days post seeding using an EVOS microscope (Thermo Fisher Scientific).

For RNA and ATAC sequencing experiments following CGRP stimulation, pancreatic epithelial cells were isolated as performed for organoid cultures until a single cell suspension was obtained. The entirety of the cell suspension was plated onto a cell culture dish with organoid media (described above) without the growth factors EGF, FGF10 and Gastrin, and supplemented with 10nM CGRP peptide or H_2_O vehicle control. Following a 2-hour stimulation, cells were collected and ice-cold MACS buffer was added 1:4 to immediately stop the stimulation by dilution. Cell suspension was centrifuged at 300g for 10 minutes at 4°C and washed three additional times with MACS buffer. To sort acinar cells, epithelial cell suspension was incubated with FcR Blocking reagent 1:100 (Miltenyi) for 20 minutes on ice and then incubated with 1:200 CD324 E-cadherin-Alexa Fluor 647 (Clone DECMA-1, #147308, Biolegend) and 1:200 *Ulex Europaeus Agglutinin I-*FITC (UEA-I-FITC, #FL-1061-2, 2BScientific), a highly specific lectin commonly used as a surface marker to identify and isolate pancreatic acinar cells (Xiao 2016), in MACS buffer for 30 minutes on ice. After three washes with MACS buffer, cells were resuspended in MACS buffer, 1 ug/mL DAPI was added and E-Cadherin positive, UEA-I positive, DAPI negative cells were fluorescent activated cell sorted (FACS) using a BD Influx cell sorter. Sorted cells were immediately centrifuged at 300g and processed for RNA and ATAC sequencing.

#### CGRP inhibition protocols

CGRP inhibition was done in vitro using the selective small molecule antagonist of the CLR/RAMP1 CGRP receptor complex, BIBN-4096 (Tocris, #4561) dissolved in DMSO. Experimental pDRG-naïve pancreatic epithelial cell co-cultures were established as above, with pDRG isolated from male animals 24 hours after the final caerulein injection. Following Matrigel set, DRG media was added supplemented with 10 μM of BIBN-4096 or DMSO control. At the time of seeding, media was additionally supplemented with Media was supplemented with 0.5 μM RhoK inhibitor Y-27632 (Stratech). Media containing BIBN-4096 or control was changed every two days, from which point Y-27632 was no longer added. To test the effect of BIBN-4096 in the growth of epithelial cells, single cell suspension of naïve pancreatic epithelial cells was obtained as above and the cells treated similarly. In both conditions, the number of organoids in each well were recorded 10 days post seeding using an EVOS microscope (Thermo Fisher Scientific) and quantified in Fiji.

#### RNA extraction

RNA extraction for acinar cells was performed with the Direct-zol RNA Microprep kit (#R2062) and for neutrophils with the QIAGEN RNeasy Micro Kit (#74004). The reason for using different due to differences in sample and starting input material, using a kit most appropriate to each cell type based on previous use. RNA extraction was performed according to manufactory instruction with in-column DNAse treatment. Acinar cell RNA concentration was measured by Qubit™ RNA High Sensitivity (Thermo, # Q10210) and submitted for in-house sequencing using Watchmaker RNA Ribo/Globin depletion library and the Novaseq 6000 sequencing platform. Neutrophil RNA concentration was measured by Bioanalyzer High Sensitivity RNA Analysis (Agilent) and submitted for in-house sequencing using NEBNExt Low Input w NEB Ultra II FS library kit and NovaSeq X sequencing platform.

#### OMNI-ATAC

ATAC sequencing of control and 2-hour CGRP stimulated acinar cells was performed using OMNI-ATAC adapted from previously published protocols (Corces 2017,Buenrostro 2015,Yoshida 2019). Detailed below.

#### Tn5 assembly

Dilution and oligo assembly of Tn5 was performed as previously described(Ma 2020,Corces 2017). Purified protein, obtained from the Francis Crick Institute protein production facilities, was diluted to 4 μM in Dilution Buffer containing 50 mM Tris, 100 mM NaCl, 0.1 mM EDTA, 1 mM DTT, 0.1% NP-40, and 50% glycerol.

For oligo annealing, two independent mixes were prepared: Mix A and Mix B. Both contained with 5 μl of 100 μM Tn5MEREV oligo, 5 μl of 100 μM of either Tn5_1 (Mix A) or Tn5_2_ME_comp (Mix B), and 40 μl of nuclease-free water. Reaction was carried out incubating each mix in a PCR thermocycler as follows: 95°C for 3 m, 65°C for 3 m and ramp down to 24°C at a rate of -1C/min. Once annealed, Mix A and Mix B were combined with 100 μl of glycerol to create a 5uM, 50% glycerol adaptor mixture.

Transposome assembly was achieved by mixing equal parts of purified diluted (4 μM) Tn5 transposase enzyme and adaptor mixture and incubate for 60 minutes at 25°C.

Oligos (5’-3’ – IDT 25nm, standard desalting):

Tn5_1 – TCGTCGGCAGCGTCAGATGTGTATAAGAGACAG Tn5_2_ME_comp – GTCTCGTGGGCTCGGAGATGTGTATAAGAGACAG TN5MEREV - /5Phos/CTGTCTCTTATACACATCT

#### Tn5 Transposition

50×10^5^ single live acinar cells (previously optimized cell number) were centrifuged at 500g for 4 minutes at 4°C and resuspended in 50 μl of ATAC resuspension buffer (RSB) (10mM Tris-HCL (Sigma) pH7.4, 10mM NaCl (Sigma), 3mM MgCl_2_ (Sigma)) with 0.1% NP40 (Thermo Fisher), 0.1% Tween-20 (Sigma) and 0.01% digitonin (Thermo Fisher) for a 4 minute incubation on ice (optimize for sorted acinar cells). 1 mL of ATAC RSB with 0.1% Tween only was added and the solution was centrifuged for at 500g 10 minutes at 4°C. ATAC samples were resuspended in 50 μl of ATAC transposase mix prepared with 25 μl of 2X Tagment DNA (TD) buffer, 2.5 μl of Tn5, 16.5 μl of 1X PBS, 0,5 μl of Digitonin 1%, 0.5 μl of 10% Tween-20 and 5 μl of nuclease free water and transposition was carried out in a thermocycler at 37°C for 30 minutes at 1000rpm. DNA was cleaned up with Zymo DNA Clean & Concentrator capped (cat# D4014, Zymo Research) according to manufacture instructions and two washes were performed, each by adding five volumes of DNA binding buffer (250 μl) followed by a centrifugation step at 10,000g for 30 seconds at room temperature, two washes with 200 μl of Wash buffer and then eluted in 21 μl of elution buffer.

#### ATAC library generation

For pre-amplification of transposed fragments and library preparation, a first 50 μl polymerase chain reaction (PCR) was performed containing 20 μl of sample DNA, 25 μl of 2X PCR Master Mix NEB (New England Biolabs, # M0541S) 1 μl of 25 μM of forward primer, 1 μl of 25 μM of reverse primer and 3 μl of water. Primer for library generation in Supplementary table 2. The PCR program was comprised of 4 steps as follows: step1 - 72 °C for 5 minutes; step2 - 98 °C for 30 seconds; step3 repeated for a total of 5 cycles - 98 °C for 10 seconds, 63 °C for 30 seconds, 72 °C for 60 seconds; step4 - 4°C hold. Amplicon was cleaned up and size selected with AMPureXP beads (#A63880, Beckman coulter) by vortexing the AMPure beads, previously allowed to reach room temperature, and adding 1.8 volumes (90 μl) to the PCR reaction. Following a 5-minute incubation, the samples were placed on an appropriate magnet and allowed to rest for 2 minutes. The liquid was discarded and the beads washed twice by adding 200 μl of freshly prepared 80% Ethanol (Sigma) tubes without disturbing the beads from the magnet and carefully removing it. Beads were let dry for 4 minutes, tubes were removed from the magnet and the beads resuspended in 18 μl of EB Buffer (Qiagen). After a 2-minute rest, the tubes were re-placed in the magnet for an additional minute to separate the beads from the clean pre-amplification PCR product.

To calculate the number of extra cycles for final sample preparation, a quantitative PCR was performed with 2 μl of sample, 10ul 2X PowerUP SYBR mix (Thermo), 2.5 μl of 25 μM forward primer, 2.5 μl of 25 μM reverse primer, and 3 μl of water using the following program: Step 1 - 98 °C for 2 minutes; Step 2 - 98 °C for 30 seconds; Step 3 - 98 °C for 10 seconds; Step 4 – 63 °C for 30 seconds; Step 5 – 72 °C for 60 seconds, repeating step 3 to 5 for a total of 20 cycles. The number of extra cycles to perform on the final sample was calculated by detrmining the number of cycles required to obtain ¼ of the maximum amplification, as described in (Buenrostro 2015).

For the final amplification, a 50 μl reaction containing 12.5 μl of sample DNA (cleaned up ATAC pre-amplified sample) 25 μl of 2X PCR Master Mix NEB (New England Biolabs, # M0541S), 2.5 μl of 25 μM forward primer and 2.5 μl of 25 μM reverse primer, was subjected to the following PCR program: Step 1 - 98 °C for 30secons ; Step 2 - 98 °C for 10 seconds; Step 3 - 63 °C for 30 seconds; Step 4 - 72 C for 60 seconds, repeating Step 2 to 4 until a total of cycles calculated previously.

The PCR product was cleaned up with AMPureXP beads (#A63880, Beckman coulter) as described above with the one difference of resuspending the beads in 30 μl water after the last bead separation ending with a 30 μl final ATAC sample.

#### Acinar cell ATAC sequencing analysis and analysis

Samples were sequenced on an Illumina NovaSeq X generating an average of 62.5 million reads per sample. Duplicate samples for Control and CGRP ATACSEQ data was analyzed using the nfcore ATCASEQ pipeline [ https://nf-co.re/atacseq/2.1.2 ]. Briefly, adapter sequences were trimmed using TrimGalore [version 0.6.7], 100bp paried end fastq reads were mapped to the mouse GRCm38 [ release 95 ] using bwa_mem [version 0.7.17]; duplicates were marked and filtered using Picard [ version 3.0.0]; Peaks were called using MACS2 (version 2.2.7.1) per replicate pair using the - -broad flag, consensus set of peaks were generated using Bedtools (version 2.30.0). Consensus peaks were annotated using the R package ChIPpeakAnno (version 3.44.0). Differential peak binding was assessed using the R package DiffBind (version 3.20.0), counting the number of reads per sample per peak using the R package subread (version 2.0.1). Pathway level enrichment of differential annotated peaks was assessed using the R package clusterProfiler (version 4.18.4) and fgsea (version 1.36.2) running R 4.5.1 within a singularity container provided by the Bioconductor project (Release 3.21). The container image was executed using Singularity (version 3.3). Data associated with the ATACSEQ experiment has been uploaded to NCBI’s Gene Expression Omnibus under the identifier GSE333849.

#### Acinar cell Bulk RNA sequencing and analysis

RNA sequencing was performed on the Illumina HiSeq 4000 system with Paired end 100bp reads. Sequencing yields were approximately 32 miilion strand specific reads per sample. Read quality trimming and adaptor removal was carried out using Trimmomatic. Reads were aligned to the mouse genome Ensembl GRCm39, [ release 89 ] using STAR [ version 2.5.2a ] and gene level counts were obtained using the RSEM package [ version 1.3.0 ]. Data was normalized, and differential expression analysis was assessed with DESeq2 [version 1.38.3 ] package within R [ 4.2.0 ], taking into account the batch variability [ ∼ batch + group ]. Genes were considered to be differentially expressed between Control and CGRP sample groups, if their adjusted p value < 0.05. Gene Set Enrichment analysis was performed for each pairwise comparison using gene lists ranked by log2FoldChange. GSEA was carried out using the fgsea package [version 1.37.4 ] with C2 and C5 gene sets. Gene signatures were considered significant if the FDR q value was less than 0.05. Data associated with the RNASEQ experiment has been uploaded to NCBI’s Gene Expression Omnibus under the identifier GSE333650.

#### Chemogenetic sensory inhibition and acute pancreatitis re-challenge

Chemogenic sensory inhibition was performed in 16-22-week-old males and females (specified in the figure legend) using R26-CAG-hM4Di^LSL^; Scn10a-Cre animals, heterozygous for both alleles. Animals were either treated with PBS (naïve cohort) or caerulein, to induce acute pancreatitis, and allowed to recover for 8 or 21 days (specified in the figure legend) before nerve inhibition was performed on AP animals only. Nerve inhibition was achieved by supplementing Clozapine N-oxide hydrochloride (25780-25mg-CAY, Cambridge Biosciences) in the drinking water for 8 days. CNO was prepared by making up a 5 mg/mL stock solution in water and using 1:1000 dilution in drinking water (5 μg/mL) corresponding to 1mg per kg of body erigh. per day. Dosages were calculated based for a typical mouse (20-27g of body weigh) which drinks approximately 5 mL of water a day, corresponding to a 25 µg of CNO daily (25 µg per 25 g mouse a day, equivalent to 1mg/kg body weigh). Following 8 consecutive days of CNO in drinking water, animals were moved to normal drinking water in the morning and subjected to a second round of acute pancreatitis 24h hours after. The 24-hour interval ensure that nerve inhibition was not sustain at the time of second AP, as reported previously(Jendryka 2019). Two days after the second AP episode, the experiment was terminated and samples collected.

#### DRG ex vivo treatments

For ex vivo treatments, pDRG explants, isolated as described above, were placed on top of a 5 μL drop of GFR Matrigel (Corning) on a 96-well plate. The DRG-containing Matrigel drop was allowed to set for 15 minutes at 37 °C and DRG media containing the treatment agents was added, details below. For all treatments, pDRGs were incubated in DRG media (described above) with the treatment of choice for 48h at 37°C. After incubations, pDRGs were washed twice in PBS and carefully removed with fine forces to a new plate. Re-plating and culture after treatment was done as described for DRG-epithelial cells co-culture. The effect of DRG treatments was assess by the number of organoids for by the co-cultured epithelial cells. Organoids cultures were recorded using EVOS microscope (Thermo Fisher Scientific) 10 days after plating and quantified in Fiji.

IL6 (mouse, #216-16-50UG, PeproTech) prepared in water and added to DRG media at a final concentration of 200 ng/mL.

G-CSF (human, Xiamen Amoytop Biotech, Lot 202109B28) was prepared in water and added to DRG media at a final concentration of 250 ng/mL.

CD45 and Neutrophil Conditioned medium was generated as described above and used 1:1 with fresh DRG media.

#### IL6 and GCSF ELISA

Detection of IL6 and G-CSF in CD45 Conditioned media was performed using the mouse IL-6 Quantikine ELISA kit (#M6000B, R&D) and the mouse G-CSF (CSF3) ELISA kit (#EMCSF3, Thermo), respectively, according to manufactory’s instructions.

Briefly, for IL6 detection, 10 μL of Calibrator Diluent RD5T were added to 40 μL of freshly isolated or snap frozen conditioned media for sample preparation (total volume 50 μL). 50 μL of Assay Diluent RD1-14 were added to the center of each well and the previously prepared 50 μL sample was added on top and the solution mixed by gentle tapping of the bottom of the plate to a flat surface. Following a overnight incubation at 4 °C, the wells were aspirated, the plate washed four times with provided 1X washing buffer in water and 100 μL of Mouse IL-6 Conjugate was added to each well for 2 hours incubation at room temperature. Wells were aspirated, washed as before and 100 μL of Substrate Solution was added to each well for a 30-minute incubation at room temperature in the dark. The reaction was stopped by 100 μL of Stop Solution and the optical density was measured using a microplate reader (TECAN) at 450nm. The amount of IL6 was determined based on standard concentration performed in paradelle.

For G-CSF detection, samples were prepared by combining 40 μL of freshly isolated or snap frozen conditioned media to 60 μL of 1X Assay diluent B. The 100 μL sample mix was added to the ELISA wells and incubated overnight at 4°C. The next day, solution was discarded, the wells were washed four times with 1X washing buffer in water and 100 µL of prepared 1X biotin conjugate was added to the wells for a 1-hour incubation at room temperature. Following an additional washing step as before, wells were incubated for 45 minutes with 100 µL of prepared Streptavidin-HRP solution at room temperature, washed four times as described and incubated with 100 µL of TMB Substrats for another 45 minutes at room temperature in the dark. The reaction was stopped by adding 50 µL of Stop Solution to each well and absorbance was measured using a microplate reader (TECAN) at 450nm. Amount of G-CSF was determined based on standard concentration performed in paradelle.

#### Flow cytometry analysis of pancreatic immune infiltrate

Profiling of immune landscape was performed by flow cytometry analsysis on pancreas isolated from male and female control and AP animals isolated 24 hours after the final caerulein injection. Single cells were obtained by mincing the pancreas using two crossed scalpel blades into small fragments and enzymatically digested in 20mL of digestion medium composed of Advanced DMEM/F12 and 1mg/mL collagenase type V from Clostridium histolyticum (Sigma-Merck). Digestion was performed for 20 minutes in a 37°C shaker at 140rpm. The reaction was stopped by adding 30mL of ice-cold MACS buffer containing (0.5% BSA (Sigma) and 250 mM EDTA (Sigma) in PBS), the samples were centrifuged for 4 min at 400xg at 4°C, and the pellet resuspended in MACS buffer and filtered twice with a 70 um and 40μm filters sequentially. The single cell suspensions were incubated with 1:100 FcR Blocking Reagent (Miltenyi Biotec) for 20 min at 4 °C, centrifuged and the pellets were incubated with fluorescently conjugated antibodies prepared in MACS (Supplementary Table 3). Each sample was stained in 300 µL of MACS to prevent acinar cell death due to high density. Cells were washed twice in MACS buffer and once in PBS. Assessment of viability was done by staining with 1:300 LIVE/DEAD™ Fixable Near IR-780 (#L34993, Invitrogen) for 15 minutes on ice. Flow cytometry analyses were performed on a BD LSR-Fortessa (BD Biosciences), BD X20 (BD Biosciences) or BD Symphony (BD Biosciences) and subsequently analysed using FlowJo v.10.4.2 (FlowJO, LCC). Gating strategy in Supplementary Table 3.

#### Neutrophil depletion

For neutrophil depletion, C57BL/6 mice were injected interperitoneally, first with rat anti-Ly6G antibody (BioXcell, clone 1A8, 50ug per mouse) in PBS or rat IgG control (Cell Sciences Unit, Francis Crick Institute) and, two hours later, with Anti-Rat Kappa Immunoglobulin Light chain (BioXCell, clone: MAR18.5, 50ug per mouse) (Boivin 2020) one day before the 3h acute pancreatitis protocol.

#### Fluorescence activated cell sorting (FACS) of neutrophils

Neutrophil cell sorting was performed as described for the flow cytometry analysis of pancreatic immune infiltrate except for the antibody combination (see Supplementary Table 3) and using 1 ug/mL DAPI for viability just before sorting. To minimise processing time and prevent activation-induced transcriptional artifacts, neutrophil isolation was performed in two sequential sorting steps. An initial low-stringency sort was used to deplete debris and enrich for neutrophils, followed by a high-purity single-cell sort to achieve the final cell population. This two-step strategy reduced the time cells spent under sorting stress while maximising yield and purity. Cell sorting was performed using BD Influx cell sorter.

#### Neutrophil RNA-seq preprocessing and gene set enrichment analysis

Male and female mice were intraperitoneally injected twice, one hour apart, with caerulein and pancreatic neutrophil were sorted 1-hour after the final caerulein injection (3-hour AP protocol) and bulk RNA sequenced. Bulk RNA-seq libraries were generated using the NEBNext Single Cell/Low Input RNA Library Prep kit and sequenced as paired-end reads on an Illumina NovaSeq X platform. Raw paired-end FASTQ files were processed in R using the packages Rfastp and Rsubread. Reads were aligned to the mouse reference genome GRCm39 using subjunc from the Rsubread package. Gene-level counts were generated using featureCounts against Ensembl GRCm39 gene annotation and differential expression analysis was performed using DESeq2.

Gene set enrichment analysis (GSEA) was performed on the complete DESeq2-ranked gene list for each contrast, without prior differential-expression thresholding. Genes were ranked by the DESeq2 Wald statistic. Ensembl gene identifiers were converted to Entrez identifiers using org.Mm.eg.db; where multiple entries mapped to the same Entrez identifier, the gene with the largest absolute ranking statistic was retained. GSEA was performed in R using clusterProfiler with the fgsea implementation. Gene sets containing 10–500 genes from KEGG, Reactome, and MSigDB Hallmark were tested. The Benjamini–Hochberg method was used to adjust for multiple testing, and pathways with adjusted P value < 0.10 were considered significant. Data associated with the RNASEQ experiment has been uploaded to NCBI’s Gene Expression Omnibus under the identifier GSE333862.

### QUANTIFICATION AND STATISTICAL ANALYSIS

Quantification of immunofluorescence on paraffin embedded samples were performed no less than 2 Levels per mouse. Unless stated otherwise values were averaged per level and then per mouse. Statistics were performed for biological replicates and number of animals specified in figure legends (n=number of animals). Specific details for each analysis are detailed below.

#### Amylase measurements by IF

Quantification of amylase reduction by immunofluorescence was performed using QuPath 0.6.0. Tissue areas were segmented into superpixels and DAPI-positive regions were excluded by determining the median DAPI intensity within each superpixel field; fields in which a nucleus was detected were omitted from downstream measurements (ROI: 2.00 µm per pixel; DAPI median threshold: >10). Amylase intensity was then quantified within the remaining superpixel fields using control samples to establish a baseline median intensity (ROI: 2.00 µm per pixel; Amylase: Median). Caerulein-treated samples were visually inspected to validate that values falling below the baseline corresponded to areas of histological damage, and a median amylase intensity threshold of 25 was established below which signal was considered reduced. The area of superpixels with a median amylase intensity below this threshold was summed and expressed as a percentage of the total measured tissue area.

#### Ki67, pH3 and STMN1 quantification

Quantification of Ki67+ and pH3+ acinar cells was performed by manual cell counting in QuPath across the entirety of three tissue levels per animal (150 µm apart). The total number of positive cells was divided by the total pancreatic tissue area to obtain a density expressed as number of cells per µm². For STMN1 quantification, a minimum of six randomly distributed regions of interest (ROIs) per tissue level were manually scored. Within each ROI, total acinar cells and STMN1+ acinar cells were counted and the proportion of STMN1+ cells was expressed as a percentage of total acinar cells.

#### pErk and βIII-Tubulin measurement

Quantification of Phospho-Erk (pErk) and βIII-Tubulin pancreatic coverage was performed in QuPath on ten randomly selected ROIs per tissue section, each covering an area of 62,500 µm². A pixel classifier was applied to the fluorescence channel at a resolution of 1.99 µm/px with a defined intensity threshold visually identified as covering the positive signal. The area classified as pErk+ or βIII-Tubulin+ was retrieved for each ROI and expressed as a percentage of the total measured tissue area.

#### pErk proximity detections

To identify cell types in close proximity to pErk+ cells, cell detection was performed in QuPath using the pErk channel as the primary detection mask. A 2 µm expansion of the cell boundary was applied to approximate the cytoplasmic compartment of neighbouring cells, and fluorescence intensity within this expanded region was used to assess the presence of the markers of interest. To distinguish true positive staining from background and staining artefacts for αSMA and βIII-Tubulin, the maximum intensity and standard deviation of each channel were plotted for all detected cells and used to determine the threshold above which staining was considered positive. pErk+ cells for which the maximum intensity and standard deviation of αSMA or βIII-Tubulin within the expanded boundary exceeded the defined threshold were classified as positive for the respective marker and expressed as a percentage of total pErk+ cells.

#### CGRP+ fibre length

Quantification of CGRP+ fibre length was performed using NeuronJ. Individual fibres were manually traced, and the cumulative length of all traced fibres was summed per image and normalised to the total pancreatic tissue area to obtain a measure of CGRP+ innervation density

#### Statistical analysis

Comparisons between two independent groups were performed using an unpaired Welch’s t-test. For matched samples derived from the same animal and subjected to different treatments ex vivo, a paired t-test was used. Comparisons across more than two groups were performed using a one-way ANOVA with appropriate post-hoc testing. For experiments measuring the extent of advanced ADM (Figures 5 and S5), as this lesion type is not uniformly distributed across the pancreatic tissue, averaging values across tissue levels per animal would obscure the spatial heterogeneity of the lesions and reduce sensitivity to detect consistent treatment effects. A two-way ANOVA was therefore performed with treatment and animal identity as factors, preserving the full within-animal variance across tissue levels while accounting for inter-animal variability and maintaining the assumption of independence between observations. For experiments including one round of pancreatitis, advanced ADM was expected to be absent under baseline conditions. Therefore, each group was independently tested against a theoretical value of zero using a one-sample t-test, to determine whether advanced ADM was significantly present.

## DATA AVAILABILITY

Data are deposited in GEO: Acinar cell bulk RNA-seq data (CGRP stimulation) GSE333650; Acinar cell ATAC-seq data (CGRP stimulation) GSE333849; Pancreatic neutrophil bulk RNA-seq data (male vs female AP) GSE333862.

## AKNOWLEDGEMENT

We thank the core facilities at the Francis Crick Institute that assisted with this work, including the Biological Research Facility (Tim Zverev, Jack Williams, Nicolas Chisholm and Ola Puchalska-Osorio), Flow Cytometry Facility (Ana Agua-Doce and Steve Lim), Experimental Histopathology Laboratory (Richard Stone, Ania Mikolajczak and Emma Nye), Light Microscopy (Matt Renshaw and Donald Bell), Advanced Sequencing Facility and Bioinformatic (James Cambell). Schematics were created with BioRender. This work was supported by the Francis Crick Institute, which receives its core funding from Cancer Research UK (CC2051-Malanchi) (CC2165-Li), the UK Medical Research Council (CC2051-Malanchi) (CC2165-Li), the Wellcome Trust (CC2051-Malanchi) (CC2165-Li), and by a Career Foundation Fellowship from Pancreatic Cancer UK (CFF2025-26_Ferreira). J.N.S. is supported by the UK Medical Research Council (MR/Y010949/1).

## AUTHORS CONTRIBUTION

R.M.M.F conceptualised and designed the study, performed most of the experiments, analysed and interpreted most of the data, and wrote the manuscript. C.B. performed the intrathecal injection of virus. E.R. analysed the neutrophil RNA sequencing data and provided technical support. A.K. assisted with OMNI-ATAC optimization and sample preparation. P.C. analysed ATAC and RNA sequencing data of CGRP treated acinar cells. E.M., M.S., S.S.T, V.B and S.R. provided input and technical support. V.B. mouse genotyping and breeding support. J.N.S. provided training on dissociation of organ-innervating DRGs and provided scientific advice. L.L. provided the Scn10a-Cre and R26-CAG-hM4Di^LSL^ mouse models and scientific advice. I.M. conceived and supervised the study, guided the interpretation of data, and assisted with the manuscript writing.

## DECLARATION OF AI-ASSISTED TECHNOLOGIES

During the preparation of this work, the author(s) used ChatGPT to shorten the text of the abstract, improve the clarity of certain paragraphs in the manuscript, and populate the resource tables based on the Materials and Methods section. The author(s) reviewed and edited the output as needed and take full responsibility for the content of the published article.

