## Supplementary Figures and Tables for "Sex-Dimorphic Neural Memory Shapes Pancreatic Tissue Resilience"

### Supplementary Figure 1

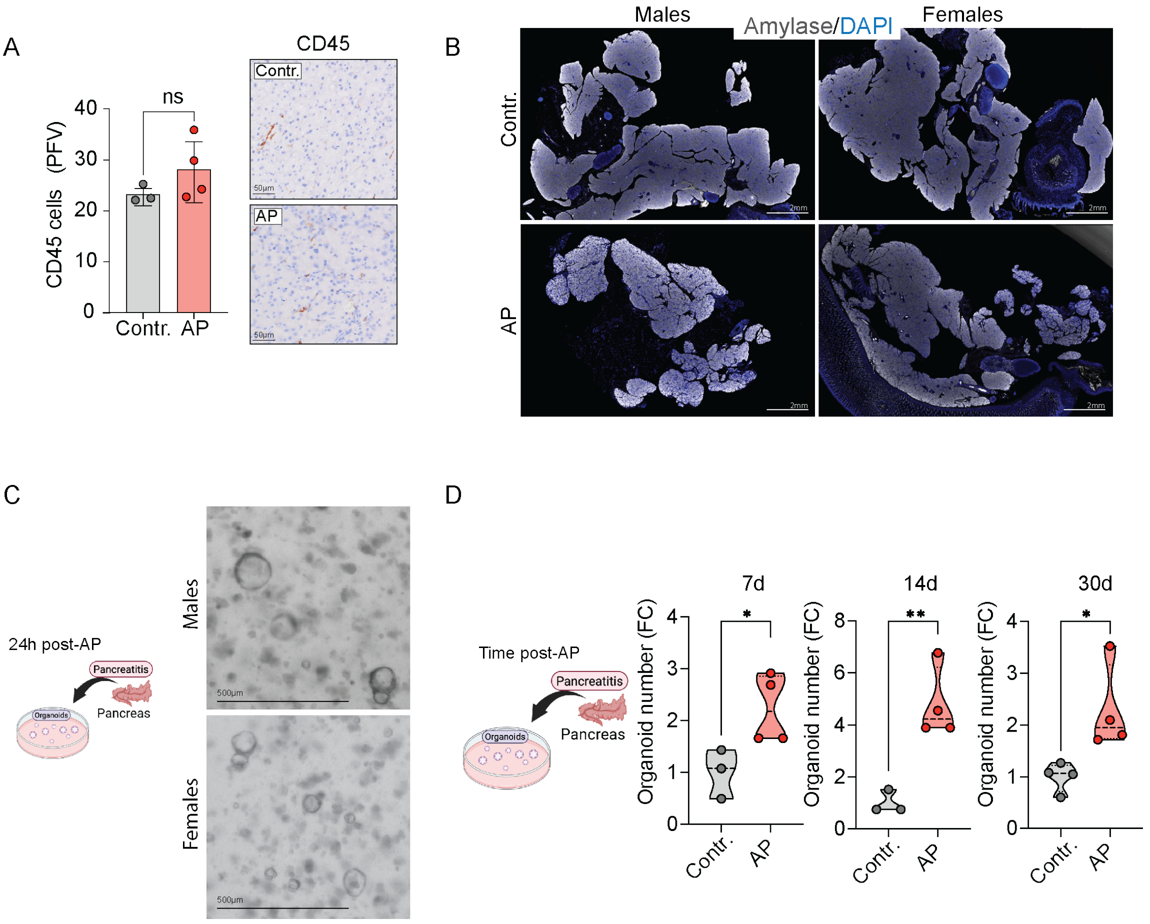


**Supplementary Figure 1 – Analysis of AP in male and female animals during injury and after repair, related to Figure 1**

**(A)** Quantification (left) of CD45 positive cells per pancreatic field of view, 7 days after final caerulein injection on male animals (control n=3, AP n=4), and representative CD45 immunohistochemistry images (Right). Data are represented as mean ± SD. Unpaired Welch's t-test. Scale bars are indicted in the figures.

**(B)** Representative amylase/DAPI immunofluorescence images of whole pancreas from male and female animals collected 24 hours after the last caerulein injection. Scale bars are indicted in the figures.

**(C)** Schematics (left) and organoid formation efficiency experiment employed to assess pancreatic epithelial plasticity and representative brightfield images (right) of organoids formed.

**(D)** Schematics (left) and organoid formation efficiency (right; number of organoids formed per well, normalised to control mean) from male pancreas at 24 hours, 14 days and 30 days after the last caerulein injection. AP performed at the same time for all groups (Male control 24h n=3, Male AP 24h n=4, Male control 14 days n=3, Male AP 14 days n=4, Male control 30 days n=4, Male AP 30 days n=4). Data are represented as median (violin plots). Unpaired Welch's t-test. Scale bars are indicted in the figures.

*p < 0.05, **p < 0.01 by Unpaired Welch's t-test.

### Supplementary Figure 2

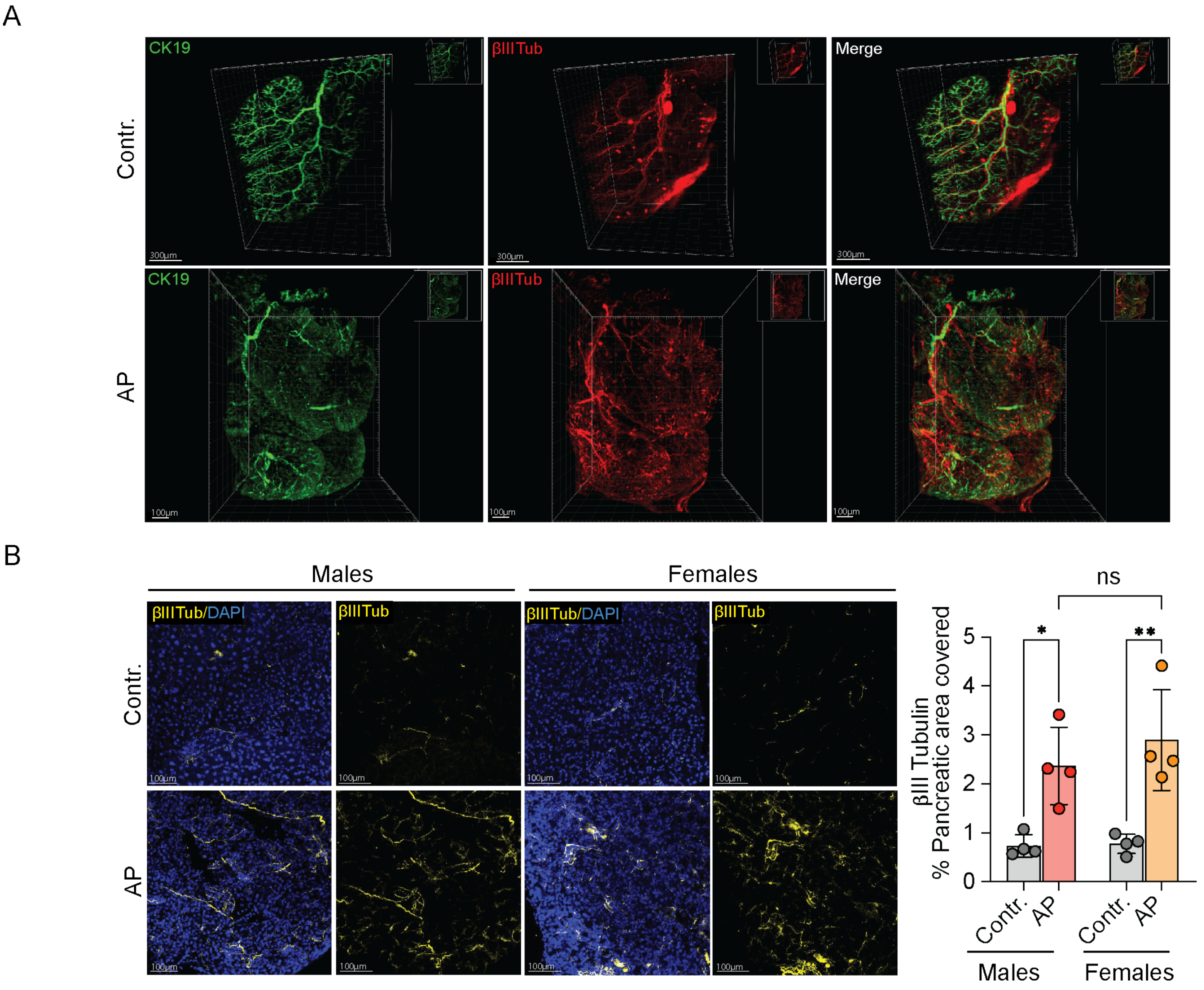


### Supplementary Figure 2 – Characterisation of AP-induced injury and repair in male and female mice, related to Figure 2

**(A)** Whole organ 3-dimensional amylase and CK19 imaging of control and male AP pancreas 24 hours after the final caerulein injection. Scale bars are indicated in the figures.

**(B)** Representative βIII-Tub/DAPI immunofluorescence images (left) of 100 μm pancreatic sections 24 hours post-AP and quantification (right) of percentage of pancreatic area covered by βIII-Tub in males (C; Male control n=4, Male AP n=4) and females (D; Female control n=4, Female AP n=4). Data are represented as mean ± SD. Unpaired Welch's t-test. Scale bars are indicted in the figures.

*$p < 0.05$ by Unpaired Welch's t-test.

### Supplementary Figure 3

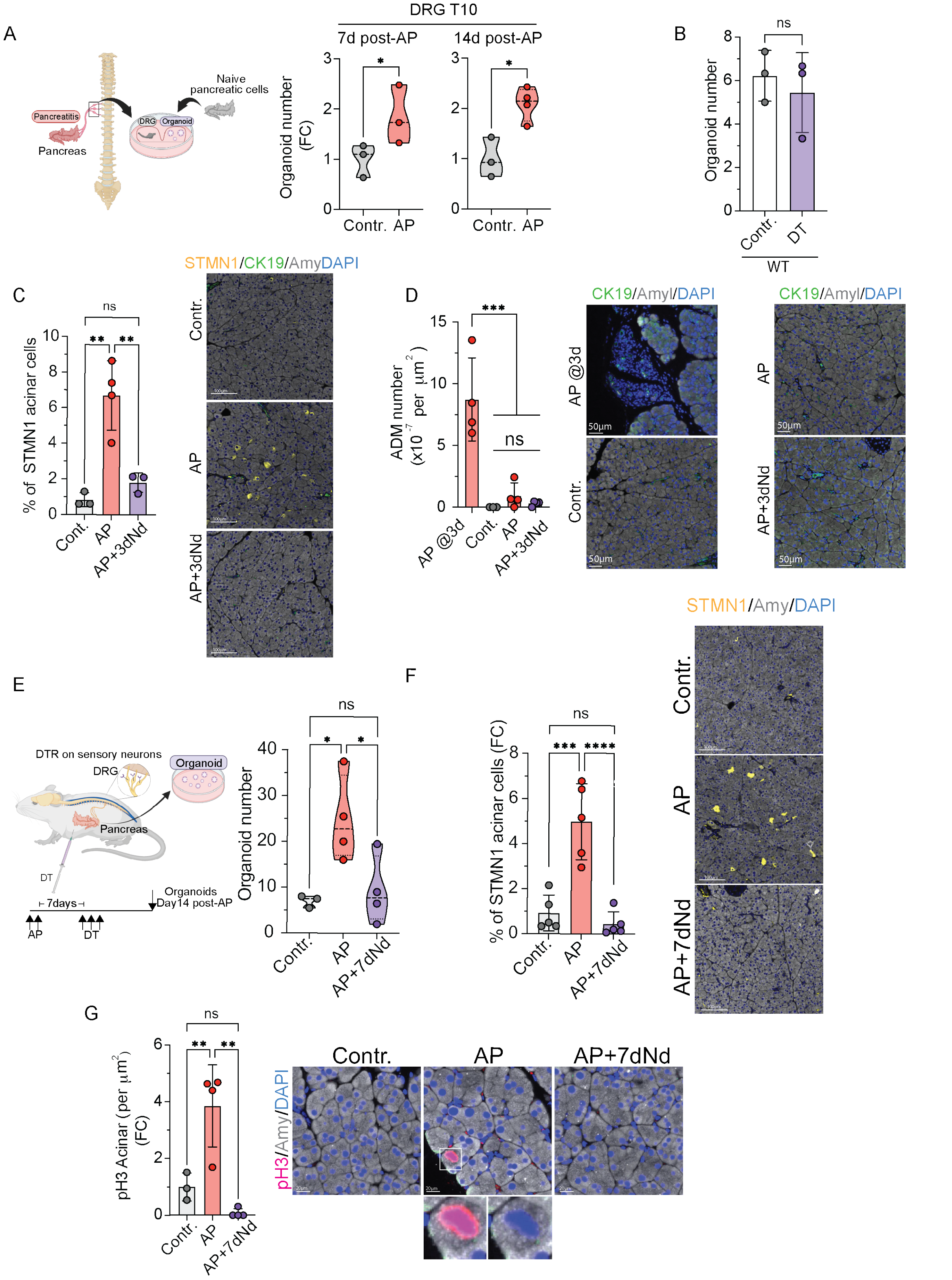


**Supplementary Figure 3 – Temporal characterisation of sensory neuron activation and nociceptor contribution to epithelial plasticity in males, related to Figure 3**

**(A)** Schematic (left) of the pDRGs and naïve epithelial cell co-culture assay for temporal characterization of neuron activation in males. Animals were subjected to AP, pDRGs were isolated 7 and 14 days after final caerulein injection and co-cultured with naïve epithelial cells in organoid formation assays. Organoid formation efficiency (right) (bottom) by timepoint (7d control n=3, 7d AP n=3, 14d control n=3, 14d AP n=4). AP was induced simultaneously across groups. Data normalised to control mean per experiment. Data are represented as median (violin plots). Unpaired Welch's t-test.

**(B)** Organoid formation efficiency of pancreatic epithelial cells isolated from control (n=3) and DT treated (n= 3) WT males to assess effect of DT treatment on epithelial plasticity.

**(C)** Quantification of percentage of SMTN1+ acinar cell (left) from early nociceptor depletion experiment in Figure 3D and representative immunofluorescence images of amylase/STMN1/CK19/DAPI. Data are represented as mean ± SD. One-way ANOVA. Scale bars are indicted in the figures.

**(D)** Quantification of number of ADM lesions in pancreatic parenchyma (left) 3 days (AP @3d n= 4), and 14 days after final caerulein in control (n= 4), AP (n=4) and nerve-depleted animals at day 3 post-AP (AP+3dNd n=4) and representative amylase/CK19/DAPI immunofluorescence images. Data are represented as mean ± SD. One-way ANOVA. Scale bars are indicted in the figures.

**(E-G)** 7 day post-AP nociceptor depletion experiment. Schematic (E, left): nociceptor neurons were depleted by DT 7 days post-AP in Scn10a-Cre;R26-DTR males, with analysis at day 14. Organoid formation efficiency (E, right; Contr. n=3, AP n=4, AP+21dNd n=4). Data normalised to control mean per experiment. Data are represented as median (violin plots); one-way ANOVA. Percentage of STMN1+ acinar cell (G, left) and representative amylase/Ki67/DAPI immunofluorescence images (F, right; Contr. n=5, AP n=5, AP+7dNd n=5). Data are represented as mean ± SD; one-way ANOVA. Quantification of pH3+ acinar cell density per $\mu m^2$ pancreatic parenchyma (G, right) and representative amylase/pH3/DAPI immunofluorescence images from the same animals. Data are represented as mean ± SD. One-way ANOVA. Scale bars are indicted in the figures.

*$p < 0.05$, **$p < 0.01$, *** $p < 0.001$, **** $p < 0.0001$ by Unpaired Welch's t-test (A-B) and one-way ANOVA (C-G)

### Supplementary Figure 4

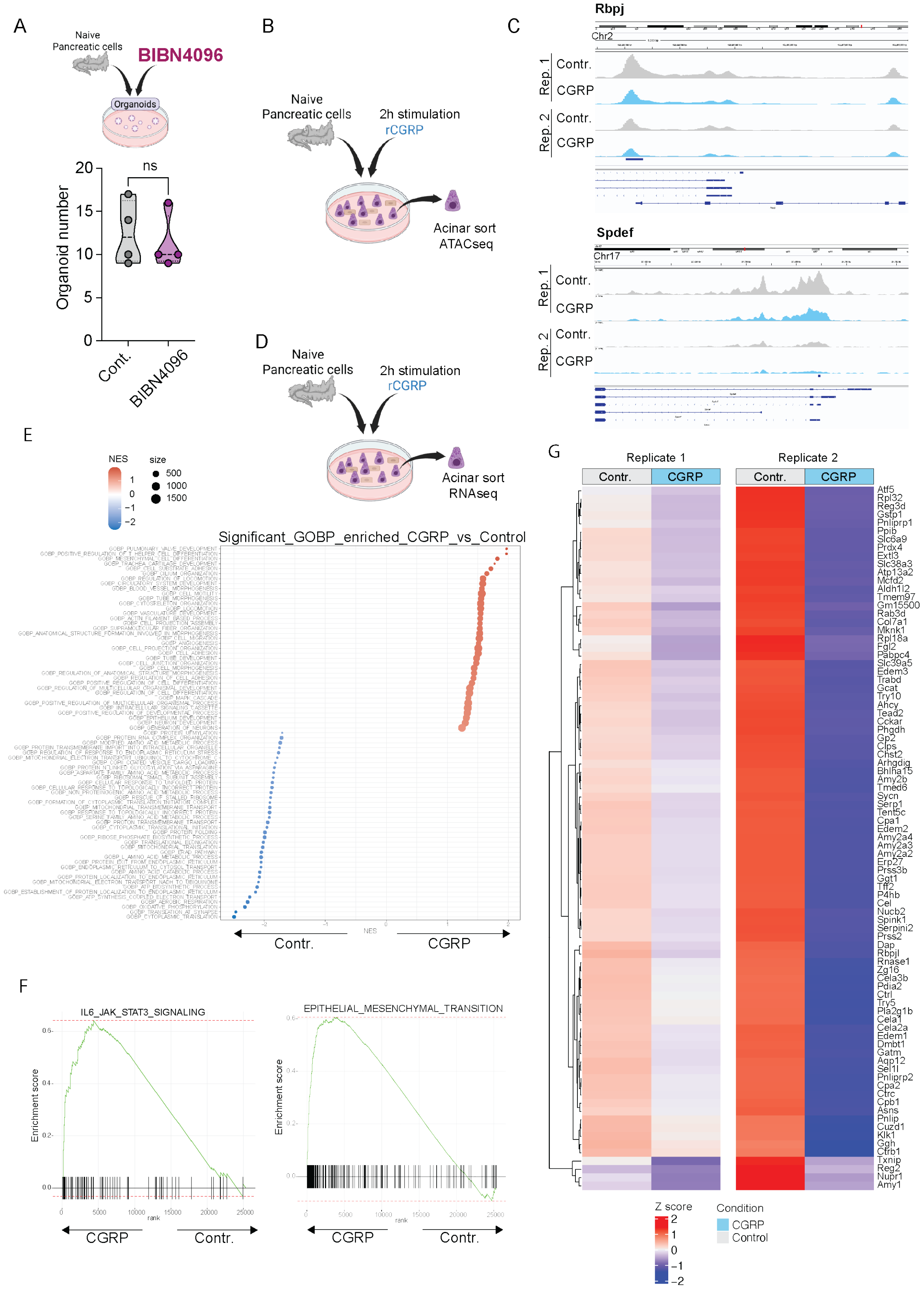


### Supplementary Figure 4 – Extended chromatin accessibility and transcriptional profiling of CGRP-stimulated acinar cells, related to Figure 4

**(A)** Schematics (top) and organoid formation efficiency (bottom; number of organoids formed per condition) of naïve pancreatic epithelial cells treated with vehicle (control n=4) and or 10 μM BIBN-4096 (BIB4096 n= 4). Data are represented as median (violin plots). Unpaired Welch's t-test

**(B-C)** Schematic (B) of 10 nM CGRP 2 hour stimulation of naïve epithelial cells followed by ATAC-sequencing of sorted acinar cells. ATAC-seq peaks (C) at Rbpj and Spdef loci.

**(D-G)** Schematic (D) of 10 nM CGRP 2 hour stimulation of naïve epithelial cells followed by RNA-sequencing of sorted acinar cells. Gene ontology biological processes (GOBP) enriched in CGRP samples (E), GSEA plots (F) showing enrichment of IL6 and EMT signalling in CGRP treated samples and 50 most deregulated gene list (G) in control and CGRP treated samples in paired replicate samples.

*$p < 0.05$ by Unpaired Welch's t-test.

**Supplementary Figure 5**

A

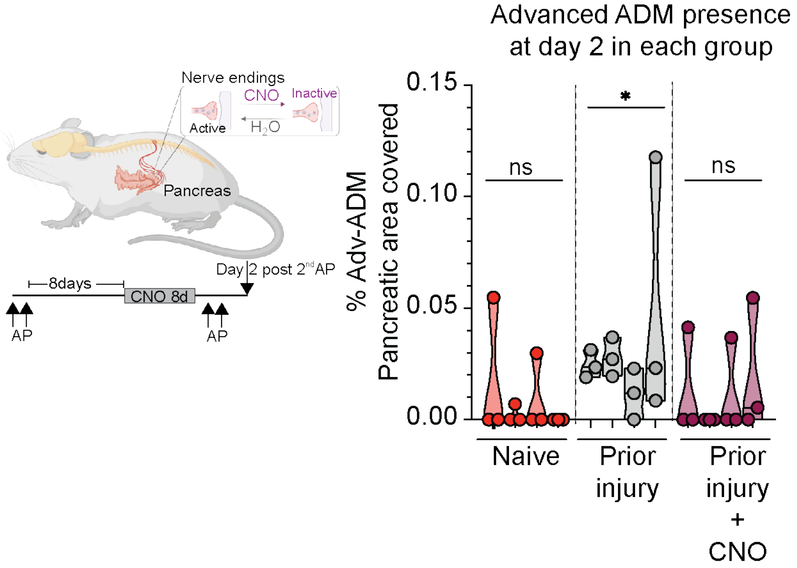


B

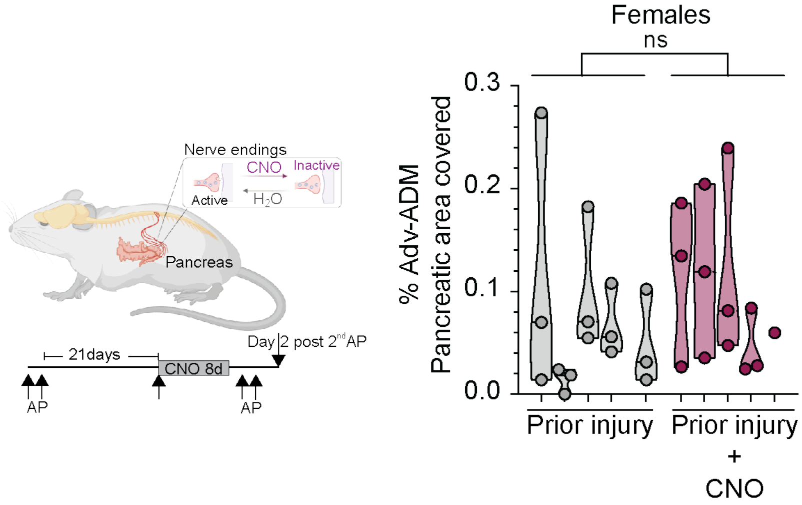


**Supplementary Figure 5 – Extended sensory inhibition analysis in males and females following AP, related to Figure 5**

**(A)** Schematic (left) of transient sensory inhibition followed by AP re-challenge. CNO, or vehicle, was delivered in drinking water for 8 days starting 8 days post-AP, prior to a second AP episode. Quantification (right) of parenchyma area covered by advanced/late ADM lesions in naïve animals (n=4) and animals with (n=5) and without (n=5) sensory inhibition. Left graph shows individual histological sections 150 µm apart (3 dots) per animal with each violin plot representing one animal. Graph on the right shows average of Advance ADM per animal. Data are represented as median (violin plots).

**(B)** Schematic (left) of transient sensory inhibition followed by AP re-challenge in females. CNO, or vehicle, was delivered in drinking water for 8 days starting 21 days post-AP, prior to a second AP episode. Quantification (right) of parenchyma area covered by advanced/late ADM lesions in animals with (n=5) and without (n=5) sensory inhibition (3 histological sections per animal, 150 µm apart). Data are represented as median (violin plots).

*$p < 0.05$ (A) where each group was tested against zero using a one-sample t-test to assess whether advanced ADM was significantly detected: n=4/group Naive (0.01825653; 0.00232206; 0.00993569; 0) Prior-Injury (0.02458025; 0.02817883; 0.01161474; 0.04976062) Prior-Injury-CNO (0.01379693; 0; 0.01224442; 0.01999644). (B) by two-way ANOVA to assess the difference in advanced ADM between the re-challenged groups.

### Supplementary Figure 6

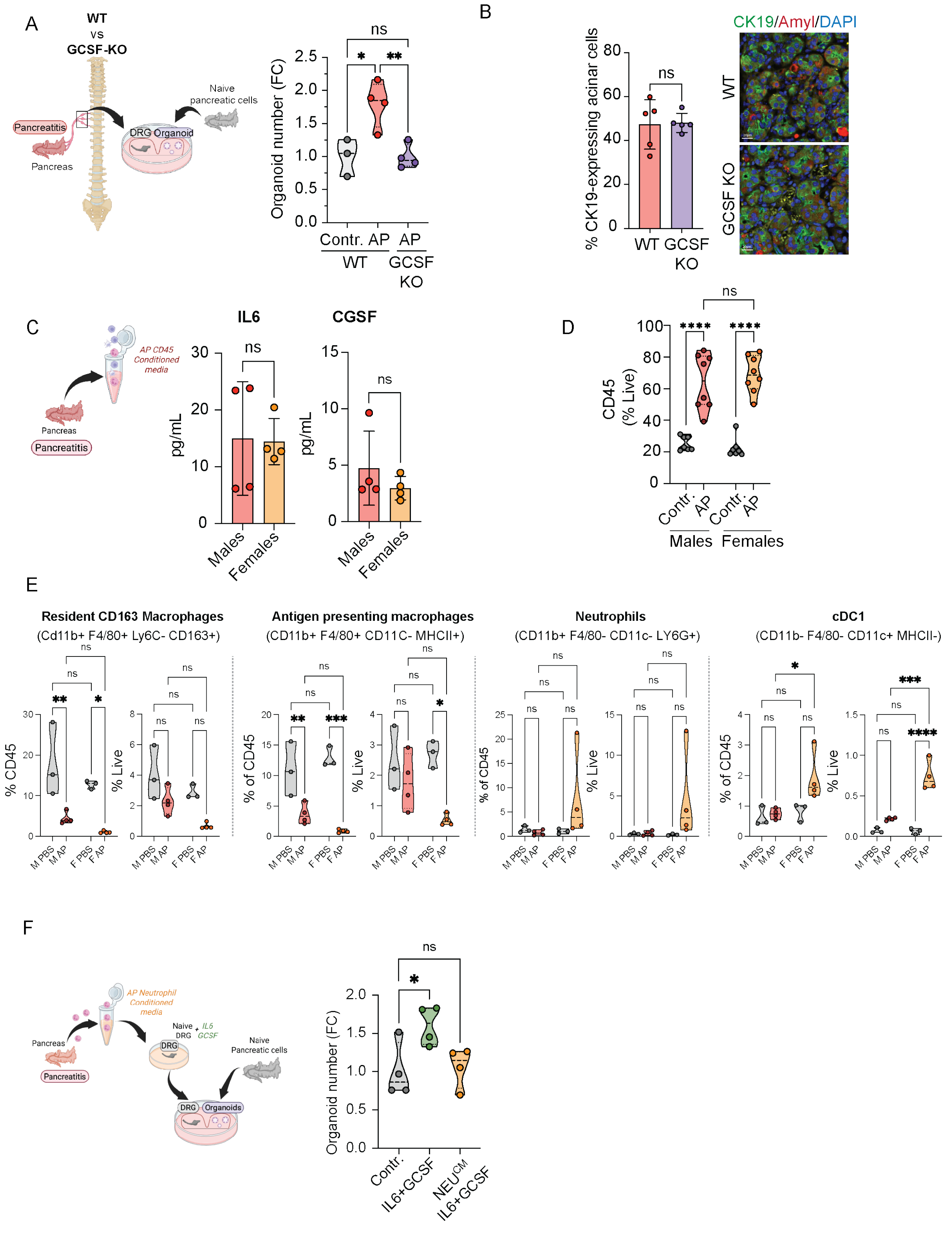


### Supplementary Figure 6 – Extended immune profiling, cytokine characterisation, and neutrophil-mediated suppression of DRG activation in AP mice, related to Figure 6

**(A)** Schematic (left) of the pDRGs-naïve epithelial cell co-culture assay to assess neuron activation and support of epithelial plasticity in WT and GCSK KO animals. pDRGs isolated

24 hours post-AP were co-cultured with naïve pancreatic epithelial cells in organoid formation condition. Organoid formation efficiency (right) by genotype (WT control n=3, AT AP n=4, GCSF-KO AP n=4). Data normalised to WT control mean. Data are represented as median (violin plots). Unpaired Welch's t-test.

**(B)** Quantification (left) of CK19+ acinar cells as a percentage of total acinar cells from WT and GCSF-KO mice 24 hours post-AP (WT n=5, Male AP n=5, and representative CK19/amylase/DAPI immunofluorescence images (right). Data are represented as mean ± SD. Unpaired Welch's t-test. Scale bars are indicted in the figures.

**(C)** Schematic (left) of conditioned media generated from male and female pancreatic-infiltrating CD45+ cells 24h after final caerulein injection and ELISA (right) to determine pg/mL of IL6 and GCSF in this media in males (n=4) and females (n=4). Data are represented as mean ± SD. Unpaired Welch's t-test.

(D) FACS-based quantification of CD45+ pancreatic-infiltrating cell populations as percentage of total live cells in control and AP male and female mice 24 hours post-injury (Male control n=7, Male AP n=8, Female control n=7, Female AP n=8). Data normalised to control mean per experiment. Data are represented as median (violin plots). One-way ANOVA.

**(E)** FACS-based quantification of individual pancreatic immune cell populations shown in the Figure 6E of the most differently present population, as a percentage of total CD45+ cells in control and AP male and female mice 24 hours post-injury (Male control n=3, Male AP n=4, Female control n=3, Female AP n=4). Data normalised to control mean. Data are represented as median (violin plots). One-way ANOVA.

**(F)** Schematic (left) of female pancreatic-infiltrating neutrophil secretome assay on DRG activation and support of epithelial plasticity. DRGs were *ex vivo* activated with 200ng/mL of IL6 and 250 ng/mL of GCSF with or without female Neutrophil CM. *Ex vivo* treated DRGs were then co-cultures with naïve epithelial cells in organoid conditions. Organoid formation efficiency (right) from epithelial cells co-cultured with naïve DRGs (control n=4), *ex vivo* activated (IL6+GCSF n=4) and *ex vivo* activated in the presence of neutrophil CM ($Neu^{CM}$ IL6+GCSF n= 4). Data normalised to control mean. Data are represented as median (violin plots). Unpaired Welch's t-test.

*p < 0.05, **p < 0.01, *** p < 0.001, **** p < 0.0001 by Unpaired Welch's t-test (B-C) and one-way ANOVA (A, D-F)

**Supplementary Table 1 -** Acinar cell identity signature

| Gene | Full Name | Function in Acinar Cell Biology | References |
|---|---|---|---|
| **CATEGORY 1: Master Transcriptional Regulators of Acinar Specification and Identity** | | | |
| Ptf1a | Pancreas transcription factor 1a | Master regulator of acinar specification and adult identity maintenance; binds ~1,000 acinar gene loci; directly activates the digestive enzyme programme; loss causes ADM and PDAC predisposition. | Hoang C.Q et al. |
| Rbpjl | Recombination signal binding protein Ig kappa J-like | Pancreas-restricted subunit of the PTF1-L complex; principal activator of digestive enzyme genes in the mature acinar cell; loss reduces acinar enzyme expression by up to 99%. | Hoang C.Q et al., Masui T et al. |
| Bhlha15 | Basic helix-loop-helix family member a15 (MIST1) | Maintains acinar polarity, cell-cell junctions and zymogen granule biogenesis; loss causes acinar disorganisation and spontaneous ADM. | Hoang C.Q et al., Pin et al. |
| Spdef | SAM pointed domain-containing ETS transcription factor | PTF1A network member; contributes to late acinar differentiation and resistance to metaplastic reprogramming. | Hoang C.Q et al. |
| Nr5a2 | Nuclear receptor subfamily 5 group A member 2 | Top-tier acinar differentiation TF; inactivation rapidly destabilises the acinar transcriptome and sensitises cells to oncogenic transformation. | Hoang C.Q et al. |
| **CATEGORY 2: Digestive Enzymes** | | | |
| Amy1 | Amylase alpha 1 | Starch-digesting enzyme; produced exclusively by acinar cells; lost upon dedifferentiation. | Hoang C.Q et al., Baron M et al. |
| Amy2a1 | Amylase alpha 2A1 | Major pancreatic amylase isoform; direct PTF1A/RBPJL target; canonical acinar identity marker. | Hoang C.Q et al., Baron M et al. |
| Amy2a4 | Amylase alpha 2A4 | Pancreatic amylase isoform; expressed in mature acinar cells; lost during ADM. | Hoang C.Q et al., Baron M et al. |
| Amy2b | Amylase alpha 2B | Pancreatic amylase isoform; canonical acinar marker; direct downstream target of the PTF1A–RBPJL complex. | Hoang C.Q et al., Azevedo-Pouly et al. |
| Try5 | Trypsinogen 5 | Trypsinogen isoform; produced exclusively by acinar cells; direct PTF1A target; premature intracellular activation initiates AP. | Hoang C.Q et al., Baron M et al. |
| Try10 | Trypsinogen 10 | Trypsinogen isoform; expressed in mature acinar cells; direct PTF1A target. | Hoang C.Q et al., Baron M et al. |
| Prss3b | Serine protease 3B (trypsinogen isoform) | Trypsinogen isoform contributing to the acinar proteolytic enzyme repertoire; downregulated in AP-induced dedifferentiation. | Hoang C.Q et al., Yang K et al. |
| Cpa1 | Carboxypeptidase A1 | Exopeptidase for C-terminal aromatic/aliphatic residues; direct PTF1A target; used as a lineage-tracing marker for acinar cells. | Hoang C.Q et al., Azevedo-Pouly et al. |
| Cpa2 | Carboxypeptidase A2 | Carboxypeptidase with preference for aromatic residues; acinar identity marker; downregulated during ADM. | Hoang C.Q et al., Azevedo-Pouly et al. |
| Cpb1 | Carboxypeptidase B1 | Exopeptidase that cleaves C-terminal basic residues; part of the acinar proteolytic enzyme repertoire. | Hoang C.Q et al., Azevedo-Pouly et al. |
| Cel | Carboxyl ester lipase | Bile salt–dependent lipase; digests cholesteryl esters and fat-soluble vitamins; directly regulated by PTF1A–RBPJL. | Hoang C.Q et al., Azevedo-Pouly et al. |
| Pnlip | Pancreatic lipase | Primary enzyme for triglyceride digestion; exclusively expressed in acinar cells; requires colipase (Clps) for activity. | Hoang C.Q et al. |
| Pnliprp1 | Pancreatic lipase related protein 1 | Lipase family member expressed in acinar cells; part of the lipid-digesting enzyme secretome. | Hoang C.Q et al., Azevedo-Pouly et al. |
| Pla2g1b | Phospholipase A2 group IB | Hydrolyses phospholipids at the sn-2 position; secreted by acinar cells; direct PTF1A target. | Hoang C.Q et al., Azevedo-Pouly et al. |
| Clps | Colipase | Essential cofactor that anchors pancreatic lipase to lipid surfaces in the presence of bile salts; exclusively expressed in acinar cells. | Hoang C.Q et al., Azevedo-Pouly et al. |
| Cela1 | Chymotrypsin-like elastase 1 | Serine protease that digests elastin and other proteins; acinar identity marker; downregulated in AP and upon dedifferentiation. | Hoang C.Q et al., Yang K et al. |
| Cela2a | Chymotrypsin-like elastase 2A | Elastolytic serine protease; part of the acinar proteolytic enzyme repertoire; downregulated in ADM. | Hoang C.Q et al., Azevedo-Pouly et al. |
| Cela3b | Chymotrypsin-like elastase 3B | Elastolytic serine protease; expressed in mature acinar cells; direct PTF1A target. | Hoang C.Q et al., Azevedo-Pouly et al. |
| Ctrc | Chymotrypsin C (caldecrin) | Regulates trypsinogen activation and degradation; marker of mature acinar cells; significantly downregulated during AP. | Hoang C.Q et al., Yang K et al. |
| Ctrb1 | Chymotrypsinogen B1 | Precursor of chymotrypsin B; synthesised exclusively by acinar cells; direct PTF1A target. | Hoang C.Q et al. |
| Spink1 | Serine peptidase inhibitor Kazal type 1 | Intracellular trypsin inhibitor produced by acinar cells; protects the cell from premature trypsin activation; mutations predispose to pancreatitis. | Hoang C.Q et al. |
| **CATEGORY 3: Zymogen Granule and Secretory Machinery** | | | |
| Gp2 | Glycoprotein 2 (zymogen granule membrane glycoprotein) | Most abundant membrane glycoprotein of the zymogen granule; GPI-anchored; accounts for ~40% of zymogen granule membrane protein; secreted with digestive enzymes into the pancreatic duct; specifically expressed by acinar cells. | Hoang C.Q et al., Yang K et al. |

| | | | |
|---|---|---|---|
| Sycn | Syncollin | Zymogen granule membrane protein; mediates granule–granule and granule–plasma membrane fusion during exocytosis; marker of the mature acinar secretory apparatus. | Hoang C.Q et al., Yang K et al. |
| Zg16 | Zymogen granule protein 16 | Lumenal component of zymogen granules; carbohydrate-binding lectin; marks mature acinar cells; downregulated during AP. | Hoang C.Q et al., Yang K et al. |
| Cuzd1 | CUB and zona pellucida-like domain 1 | Zymogen granule membrane component; specifically expressed in mature acinar cells; downregulated during AP-induced dedifferentiation. | Yang K et al. |
| **CATEGORY 4: ER Quality Control and Protein Folding Machinery** | | | |
| Pdia2 | Protein disulfide isomerase A2 | Pancreas-specific PDI isoform; catalyses disulfide bond formation and isomerisation during folding of secretory proteins; part of the specialised acinar ER quality control apparatus. | Hoang C.Q et al. |
| Erp27 | Endoplasmic reticulum protein 27 | ER-resident disulfide isomerase family member; assists folding of digestive enzyme precursors; part of the acinar ER quality control machinery. | Hoang C.Q et al. |
| P4hb | Prolyl 4-hydroxylase subunit beta (PDI) | Major ER protein disulfide isomerase; essential for correct folding of all acinar secretory proteins; part of the highly attuned acinar ER surveillance system maintained by PTF1A. | Hoang C.Q et al. |
| Serp1 | Stress-associated ER protein 1 | Stabilises stalled translocon-associated ribosomes under ER stress; part of the acinar machinery managing the extreme burden of secretory protein synthesis. | Hoang C.Q et al. |
| Edem1 | ER degradation-enhancing alpha-mannosidase-like protein 1 | ERAD component; recognises and targets misfolded glycoproteins for proteasomal degradation; essential for maintaining ER homeostasis in acinar cells. | Hoang C.Q et al., Yang K et al. |
| Edem2 | ER degradation-enhancing alpha-mannosidase-like protein 2 | ERAD component; works in parallel with EDEM1 to trim mannose residues on misfolded glycoproteins; part of the acinar ER quality control programme. | Hoang C.Q et al., Yang K et al. |
| Tent5c | Terminal nucleotidyltransferase 5C | Cytoplasmic poly(A) polymerase enriched in secretory cells; stabilises mRNAs encoding digestive enzyme precursors by extending their poly(A) tails; part of the acinar post-transcriptional regulatory machinery. | Hoang C.Q et al. |
| **CATEGORY 5: Metabolic Support and Other Functional Markers** | | | |
| Aldob | Aldolase B | Glycolytic enzyme; supports the potent anabolic metabolism required to sustain the massive protein synthesis demands of the mature acinar cell; directly maintained by the PTF1A transcriptional programme. | Hoang C.Q et al. |
| Anpep | Aminopeptidase N (CD13) | Cell-surface metalloprotease expressed in acinar cells; involved in peptide processing at the luminal surface of the exocrine pancreas. | Hoang C.Q et al. |
| Tc2n | Tandem C2 domains nuclear protein | Calcium sensor involved in regulated membrane fusion; expressed in acinar cells where $Ca^{2+}$-dependent exocytosis of zymogen granules is the primary secretory mechanism. | Hoang C.Q et al. |
| Tmem97 | Transmembrane protein 97 (MAC30) | Integral membrane protein expressed in acinar cells; associated with intracellular membrane organisation supporting the high-throughput secretory activity of the mature acinar cell. | Hoang C.Q et al. |


| Abbreviation | Full Citation |
|---|---|
| Hoang C.Q et al. | Hoang C.Q et al. Transcriptional Maintenance of Pancreatic Acinar Identity, Differentiation, and Homeostasis by PTF1A. Mol Cell Biol. 2016;36(24):3033–3047. |
| Azevedo-Pouly et al. | Azevedo-Pouly et al. Key transcriptional effectors of the pancreatic acinar phenotype and oncogenic transformation. PLoS ONE. 2023, 18(10):e0291512 |
| Masui T et al. | Masui T et al. Replacement of Rbpj with Rbpjl in the PTF1 complex controls the final maturation of pancreatic acinar cells. Gastroenterology. 2010;139(1):270-280. |
| Baron M et al. | Baron M et al. A Single-Cell Transcriptomic Map of the Human and Mouse Pancreas Reveals Inter- and Intra-Cell Population Structure. Cell Systems. 2016;3(4):346-360. |
| Zhu et al. 2024 | Yang K et al. The integration of single-cell and bulk RNA-seq atlas reveals ERS-mediated acinar cell damage in acute pancreatitis. J Transl Med. 2024, 22(1):346 |
| Pin et al. 2001 | Pin CL et al. The bHLH transcription factor Mist1 is required to maintain exocrine pancreas cell organization and acinar cell identity. J Cell Biol. 2001;155(4):519-530. |

### **Supplementary Table 2** - ATAC library

| *Sample* | *Primer pair* |
|---|---|
| Q_2h_2 | NXT N703-N505 (TTCTGCCT-GTAAGGAG) |
| CGRP_10nM_: | NXT N704-N505 (GCTCAGGA-GTAAGGAG) |
| Q_2h_3 | NXT N705-N505 (AGGAGTCC-GTAAGGAG) |
| CGRP_10nM_: | NXT N706-N505 (CATGCCTA-GTAAGGAG) |

| *Primer name* | *Index (Nextera)* | *Sequence 5'-3'* |
|---|---|---|
| Ad1.3_N703_F | TTCTGCCT | CAAGCAGAAGACGGCATACGAGATTTCTGCCTGTCTCGTGGGCTCGGAGATGT |
| Ad1.4_N704_F | GCTCAGGA | CAAGCAGAAGACGGCATACGAGATGCTCAGGAGTCTCGTGGGCTCGGAGATGT |
| Ad1.5_N705_F | AGGAGTCC | CAAGCAGAAGACGGCATACGAGATAGGAGTCCGTCTCGTGGGCTCGGAGATGT |
| Ad1.6_N706_F | CATGCCTA | CAAGCAGAAGACGGCATACGAGATCATGCCTAGTCTCGTGGGCTCGGAGATGT |
| Ad2.5_N505_F | GTAAGGAG | AATGATACGGCGACCACCGAGATCTACACGTAAGGAGTCGTCGGCAGCGTCAGATGTG |

**Supplementary Table 3 -** FACS antibodies and gating strategies

| *Antigen* | *Clone* | *Fluorophore* | *Dilution* | *Catalogue number* | *Supplier* |
|---|---|---|---|---|---|
| CD3 | 17A2 | APC/Cy7 | 1:200 | 100222 | BioLegend |
| B220 | RA3-6B2 | APC/Cy7 | 1:200 | 103224 | BioLegend |
| CD45 | 30-F11 | BUV395 | 1:200 | 564279 | BD Bioscience |
| CD11b | M1/70 | FITC | 1:200 | 101206 | BioLegend |
| CD11c | HL3 | BUV737 | 1:200 | 612796 | BD Bioscience |
| Ly6G | 1A8 | BV605 | 1:200 | 127639 | BioLegend |
| Ly6C | HK1.4 | BB700 | 1:200 | 755203 | BD Optibuild |
| NK1.1 | APC | PK136 | 1:200 | 108705 | BioLegend |
| CD163 | PE | TNKUPJ | 1:200 | 12-1631-80 | eBioscience |
| F4/80 | efluor450 | BM8 | 1:200 | 48-4801-82 | eBioscience |
| MHCII | BV711 | M5/114 | 1:200 | 107643 | BioLegend |

| **Near-IR for viability** | | | | | | |
|---|---|---|---|---|---|---|
| Live CD45+ | -> | CD11b+ F4/80- | -> | Ly6C+CD163+ **Resident CD163 macrophages** | | |
| | | | -> | Ly6C+CD163- **Recruited Inflammatory macrophages** | | |
| | | | -> | Ly6C-CD163+ **Recruited transitioning macrophages** | | |
| | | | -> | Ly6C-CD163- | -> | CD11c+ MHCII+ **Mo-derived macrophages DC-like** |
| | | | | | -> | CD11c- MHCII+ **Antigen Presenting macrophages** |
| | | | | | -> | CD11c- MHCII- **Other macrophages** |
| | -> | CD11b+ F4/80- | -> | Ly6G+ **Neutrophils** | | |
| | | | -> | Ly6G- | -> | Ly6C+ MHCII- **Classic eraly inflammatory monocytes** |
| | | | | | -> | Ly6C+ MHCII+ **Transitioning DC monocytes** |
| | | | | | -> | Ly6C- MHCII+ **MoDC** |
| | -> | CD11b- F4/80- | -> | CD11c+ NK1.1+ **Activated NK** | | |
| | | | -> | CD11c- NK1.1+ **NK** | | |
| | | | -> | CD11c- NK1.1- **ILCs** | | |
| | | | -> | CD163-MHCII- **cDC1** | | |
| | | | -> | CD163-MHCII- p**DC** | | |

| *Antigen* | *Clone* | *Fluorophore* | *Dilution* | *Catalogue number* | *Supplier* |
|---|---|---|---|---|---|
| Ly6G | 1A8 | PE | 1:200 | 551461 | BD Pharmigen |
| CD11b | M1/70 | APC | 1:200 | 101212 | Biolegend |
